# Single cell genomics reveals plastid-lacking Picozoa are close relatives of red algae

**DOI:** 10.1101/2021.04.14.439778

**Authors:** Max E. Schön, Vasily V. Zlatogursky, Rohan P. Singh, Camille Poirier, Susanne Wilken, Varsha Mathur, Jürgen F. H. Strassert, Jarone Pinhassi, Alexandra Z. Worden, Patrick J. Keeling, Thijs J. G. Ettema, Jeremy G. Wideman, Fabien Burki

## Abstract

The endosymbiotic origin of plastids from cyanobacteria gave eukaryotes photosynthetic capabilities and launched the diversification of countless forms of algae. These primary plastids are found in members of the eukaryotic supergroup Archaeplastida. All known archaeplastids still retain some form of primary plastids, which are widely assumed to have a single origin. Here, we used single-cell genomics from natural samples combined with phylogenomics to infer the evolutionary origin of the phylum Picozoa, a globally distributed but seemingly rare group of marine microbial heterotrophic eukaryotes. Strikingly, the analysis of 43 single-cell genomes shows that Picozoa belong to Archaeplastida, specifically related to red algae and the phagotrophic rhodelphids. These picozoan genomes support the hypothesis that Picozoa lack a plastid, and further reveal no evidence of an early cryptic endosymbiosis with cyanobacteria. These findings change our understanding of plastid evolution as they either represent the first complete plastid loss in a free-living taxon, or indicate that red algae and rhodelphids obtained their plastids independently of other archaeplastids.

## Introduction

The origin of plastids by endosymbiosis between a eukaryotic host and cyanobacteria was a fundamental transition in eukaryotic evolution, giving rise to the first photosynthetic eukaryotes. These ancient primary plastids, estimated to have originated >1.8 billion years ago^1^, are found in Rhodophyta (red algae), Chloroplastida (green algae, including land plants), and Glaucophyta (glaucophytes)—together forming the eukaryotic supergroup Archaeplastida^2^. Unravelling the complex sequence of events leading to the establishment of the cyanobacterial endosymbiont in Archaeplastida is complicated by antiquity, and by the current lack of modern descendants of early-diverging relatives of the main archaeplastidan groups in culture collections or sequence databases. Indeed, the only other known example of primary endosymbiosis are the chromatophores in one unrelated genus of amoeba (*Paulinella*), which originated about a billion years later^3,4^. Recently, two newly described phyla (Prasinodermophyta and Rhodelphidia) were found to branch as sister to green and red algae, respectively^5,6^. Most transformative was the discovery that rhodelphids are obligate phagotrophs that maintain cryptic non-photosynthetic plastids, implying that the ancestor of red algae was mixotrophic, a finding that greatly alters our perspectives on early archaeplastid evolution^5^.

While there is substantial evidence that Archaeplastida is a group descended from a photosynthetic ancestor, non-photosynthetic and plastid-lacking lineages have been found to branch near the base or even within archaeplastids in phylogenomic trees. For example, Cryptista (which includes plastid-lacking and secondary plastid-containing species), have been inferred to be sister to either green algae and glaucophytes^7^, or red algae^5,8^, although other phylogenomic analyses have recovered the monophyly of Archaeplastida to the exclusion of the cryptists^5,9,10^. Another non-photosynthetic group that recently showed affinities to red algae based on phylogenomics is Picozoa^5,9,10^. But as for cryptists, the position of Picozoa has lacked consistent support, mostly because there is no member of Picozoa available in continuous culture, and genomic data are currently restricted to a few, incomplete, single amplified genomes (SAGs)^11^. Thus, the origin of Picozoa remains unclear.

Picozoa (previously known as picobiliphytes) were first described in 2007 in marine environmental clone libraries of the 18S ribosomal RNA (rRNA) gene and observed by epifluorescence microscopy in temperate waters^12^. Due to orange autofluorescence reminiscent of the photosynthetic pigment phycobiliprotein and emanating from an organelle-like structure, picozoans were initially described as likely containing a plastid. Orange fluorescence was also observed in association with these uncultured cells in subtropical waters^13^. However, the hypothesis that the cells were photosynthetic was challenged by the characterization of SAG data from three picozoan cells isolated by fluorescence-activated cell sorting (FACS)^11^. The analysis of these SAGs revealed neither plastid DNA nor nuclear-encoded plastid-targeted proteins, but the scope of these conclusions is limited due to the small number of analyzed cells and their highly fragmented and incomplete genomes^11^. Most interestingly, a transient culture was later established, enabling the formal description of the first (and so far only) picozoan species–*Picomonas judraskeda*–as well as ultrastructural observations with electron microscopy^14^. These observations revealed an unusual structural feature in two body parts, a feeding strategy by endocytosis of nano-sized colloid particles, and confirmed the absence of plastids^14^. Only the 18S rRNA gene sequence of *P. judraskeda* is available as the transient culture was lost before genomic data could be generated.

Here, we present an analysis of genomic data from 43 picozoan single-cell genomes sorted with FACS from the Pacific Ocean off the California coast and from the Baltic Sea. Using a gene and taxon-rich phylogenomic dataset, these new data allowed us to robustly infer Picozoa as a lineage of archaeplastids, branching with red algae and rhodelphids. With this expanded genomic dataset, we confirm Picozoa as the first archaeplastid lineage lacking a plastid. We discuss the important implications that these results have on our understanding of the origin of plastids.

## Results

### Single cell assembled genomes representative of Picozoa diversity

We isolated 43 picozoan cells (40 from the eastern North Pacific off the coast of California, 3 from the Baltic Sea) using FACS and performed whole genome amplification by multiple displacement amplification (MDA). The taxonomic affiliation of the SAGs was determined either by PCR with Picozoa-specific primers^14^ or 18S rRNA gene sequencing using general eukaryotic primers, followed by Illumina sequencing of the MDA products (see methods). The sequencing reads were assembled into genomic contigs, with a total assembly size ranging from 350 kbp to 66 Mbp (Fig 1a, Table S1). From these contigs, the 18S rRNA gene was found in 37 out of the 43 SAGs, which we used to build a phylogenetic tree with reference sequences from the protist ribosomal reference PR2 database (Fig S1). Based on this tree, we identified 6 groups representing 32 SAGs that possessed nearly identical 18S rRNA gene sequences. These SAGs with identical ribotype were reassembled by pooling all reads in order to obtain longer, more complete co-assemblies (CO-SAGs). The genome size of the CO-SAGs ranged from 32 Mbp to 109 Mbp (Fig 1a, Tab S1), an increase of 5% to 45% over individual SAGs. The genome completeness of the SAGs and CO-SAGs was estimated based on two datasets: (i) a set of 255 eukaryotic marker genes available in BUSCO^15^, and (ii) a set of 317 conserved marker genes derived from a previous pan-eukaryote phylogenomic dataset^1^ that we used here as starting point (Fig 1b). These comparisons showed that while most SAGs were highly incomplete (Fig 1a and b), the CO-SAGs were generally more complete (up to 60%). When taken together, 90% of the BUSCO markers and 88% of the phylogenomic markers were present in at least one assembly, suggesting that while the single-cell genome assemblies are fragmentary, they together represent a much more complete Picozoa meta-assembly.

**Figure 1.**
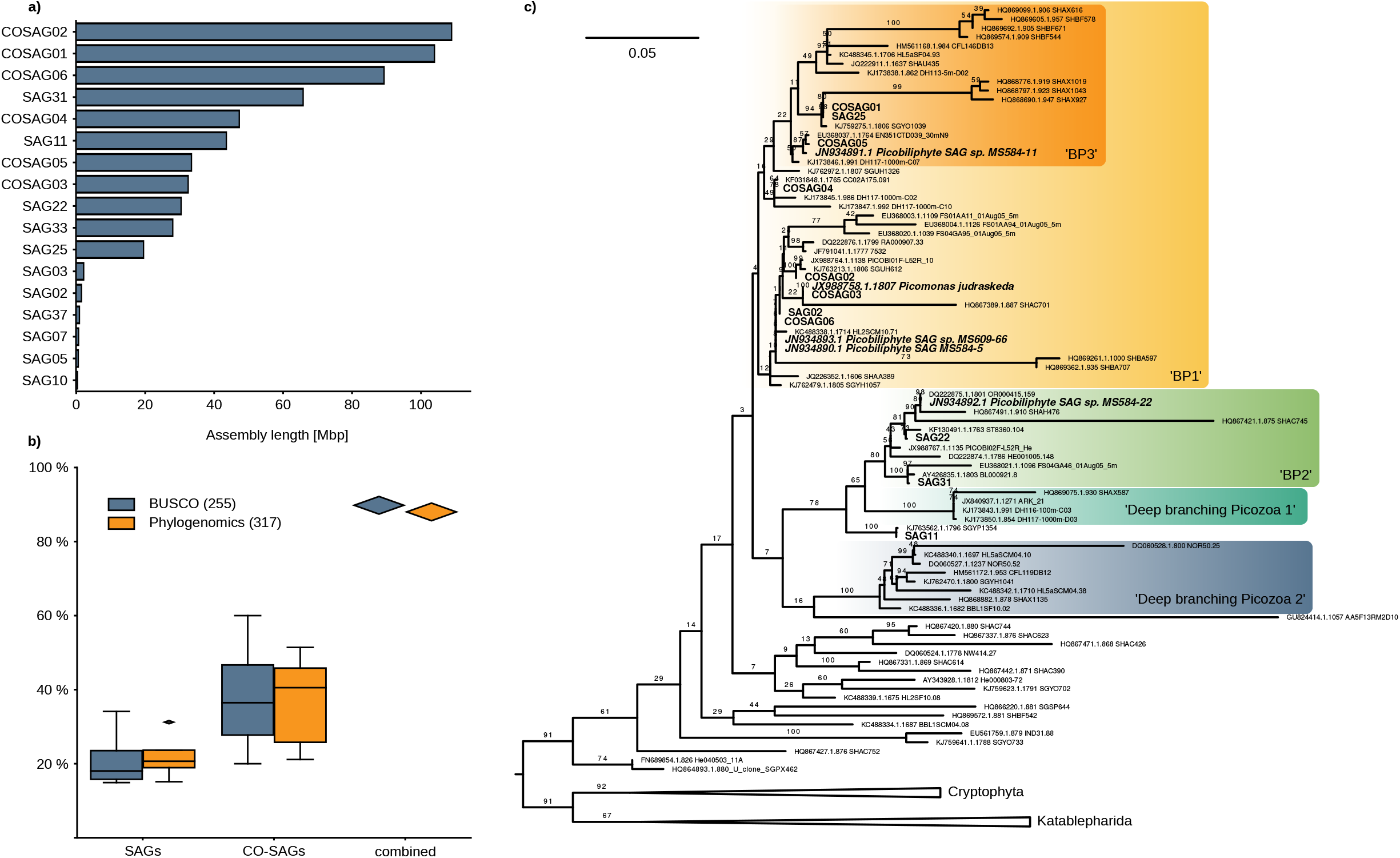
**a)** Assembly length in Mbp for 17 SAGs and CO-SAGs used for further analysis. **b)** Estimated completeness of the 10 most complete SAGs and CO-SAGs as assessed using presence/absence of the BUSCO dataset of 255 eukaryotic markers and a dataset of 317 Phylogenomic marker genes. These 10 assemblies were used for the phylogenomic inference. **c)** Maximum likelihood tree of the 18S rRNA gene, reconstructed using the model GTR+R4+F while support was estimated with 100 non-parametric bootstrap replicates in IQ-TREE. Picozoa CO-SAGs and SAGs are written in bold, the sequences of *Picomonas judraskeda* and the SAGs from Yoon et al. (2011)^11^ in bold italic. The group labels BP1-3 are from Cuvelier et al. (2008)^13^ and deep branching lineages from Moreira and López-Garcia (2014)^16^.

The final 17 assemblies (11 SAGs and 6 CO-SAGs) were mainly placed within the three proposed groups of Picozoa BP1-3 (Fig 1c), sensu Cuvelier et al. (2008)^13^. One SAG (SAG11) was placed outside of these groups. The deep-branching picozoan lineages identified by Moreira and López-García (2014)^16^, as well as other possibly early-diverging lineages were not represented in our data (Fig 1c). Interestingly, one CO-SAG (COSAG03) was closely related (18S rRNA gene 100% identical) to the only described species, *Picomonas judraskeda*, for which no genomic data are available. Using our assemblies and reference sequences from PR2 as queries, we identified by sequence identity 362 OTUs related to Picozoa (≥ 90 %) in the data provided by the *Tara* Oceans project^17^. Picozoa were found in all major oceanic regions, but had generally low relative abundance in V9 18S rRNA gene amplicon data (less than 1% of the eukaryotic fractions in most cases, Fig S2). An exception was the Southern Ocean between South America and Antarctica, where the Picozoa-related OTUs in one sample represented up to 30% of the V9 18S rRNA gene amplicons. Thus, Picozoa seems widespread in the oceans but generally low in abundance based on available sampling, although they can reach higher relative abundances in at least circumpolar waters.

### Phylogenomic dataset construction

To infer the evolutionary origin of Picozoa, we expanded on a phylogenomic dataset that contained a broad sampling of eukaryotes and a large number of genes that was recently used to study deep nodes in the eukaryotic tree^1^. Homologues from the SAGs and CO-SAGs as well as a number of newly sequenced key eukaryotes were added to each single gene (see Table S2 for a list of taxa). After careful examination of the single genes for contamination and orthology based on individual phylogenies (see material and methods), we retained all six CO-SAGs and four individual SAGs together with the available SAG MS584-11 from a previous study^11^. The rest of the SAGs were excluded due to poor data coverage (less than 5 markers present) and, in one case (SAG33), because it was heavily contaminated with sequences from a cryptophyte (see Data availability for access to the gene trees). In total, our phylogenomic dataset contained 317 protein-coding genes, with orthologues from Picozoa included in 279 genes (88%), and 794 taxa (Fig 1b). This represents an increase in gene coverage from 18% to 88% compared to the previously available genomic data for Picozoa. The most complete assembly was CO-SAG01, from which we identified orthologues for 163 (51%) of the markers.

### Picozoa group with Rhodophyta and Rhodelphidia

Concatenated protein alignments of the curated 317 genes were used to infer the phylogenetic placement of Picozoa in the eukaryotic Tree of Life. Initially, a maximum likelihood (ML) tree was reconstructed from the complete 794-taxa dataset using the site-homogeneous model LG+F+G and ultrafast bootstrap support with 1,000 replicates (Fig S3). This analysis placed Picozoa together with a clade comprising red algae and rhodelphids with strong support (100% UFBoot2), but the monophyly of Archaeplastida was not recovered due to the internal placement of the cryptists. To further investigate the position of Picozoa, we applied better-fitting site-heterogeneous models to a reduced dataset of 67 taxa, since these models are computationally much more demanding. The process of taxon reduction was driven by the requirement of maintaining representation from all major groups, while focusing sampling on the part of the tree where Picozoa most likely belong to, i.e. Archaeplastida, TSAR, Haptista and Cryptista. We also merged several closely related lineages into OTUs based on the initial ML tree in order to reduce missing data (Table S3). This 67-taxa dataset was used in ML and Bayesian analyses with the best-fitting site-heterogeneous models LG+C60+F+G+PMSF (with non-parametric bootstrapping) and CAT+GTR+G, respectively. Both ML and Bayesian analyses produced highly similar trees, and received maximal support for the majority of relationships, including deep divergences (Fig 2). Most interestingly, both analyses recovered the monophyly of Archaeplastida (BS=93%; PP=1), with cryptists as sister lineage (BS=100%; PP=1). Consistent with the initial ML tree (Fig S3), red algae and rhodelphids branched together (BS=95%; PP=1), with Picozoa as their sister with full support (BS=100%; PP=1). This grouping was robust to fast-evolving sites removal analysis (Fig S4), trimming of the 25% and 50% compositionally most biased sites (Fig S5), and was also recovered in a supertree method (ASTRAL-III) consistent with the multi-species coalescent model (Fig S6). Although this group is robust, we observed one variation in the branching order between Picozoa, rhodelphids and red algae when trimming the 50% most heterogenous sites (Fig S7) and after removing genes with less than two picozoan sequences (Fig S8). In these analyses, Picozoa and red algae were most closely related, although this relationship was never significantly supported. An approximately unbiased (AU) test rejected all tested topologies except in the two cases where Picozoa branched as the closest sister to red algae (p=0.237) and the topology of Fig 2 (p=0.822; Table S4). Finally, we identified in Picozoa and rhodelphids a two amino acids replacement signature in the eukaryotic translation elongation factor 2 protein (SA instead of the ancestral GS residues, see Supplementary Data S1) that was previously shown to unite red and green algae (and land plants), haptophytes and some cryptists^18^. The presence of SA in Picozoa supports their affiliation with red algae and rhodelphids.

**Figure 2.**
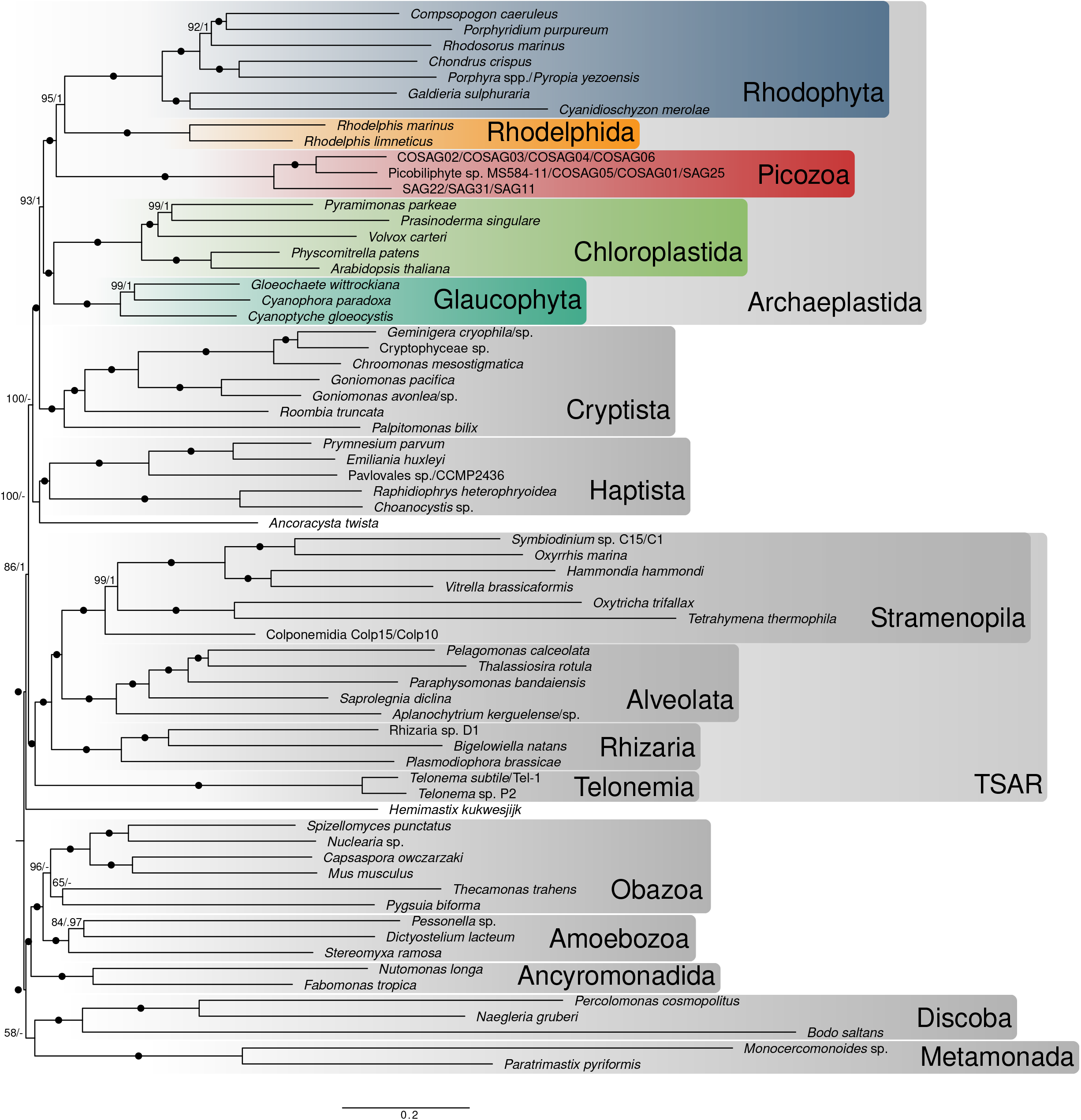
Maximum likelihood tree of eukaryotic species showing the position of Picozoa. The tree is based on the concatenated alignment of 317 marker genes and was reconstructed using the site-heterogeneous model LG+C60+F+G-PMSF. Support values correspond to 100 non-parametric bootstrap replicates/ posterior probability values estimated using PhyloBayes CAT-GTR-G. Black circles denote full support (=100/1.0).

### Picozoa SAGs show no evidence of a plastid

Since there have been conflicting conclusions about the occurrence of plastids in picozoans, we extensively searched our genomic data for evidence of cryptic plastids. First, we searched the SAG and CO-SAG assemblies for plastidial contigs as evidence of a plastid genome. While there were some contigs that initially showed similarities to reference plastid genomes, these were all rejected as bacterial (non-cyanobacterial) contamination upon closer inspection. In contrast, mitochondrial contigs were readily identified in 26 of 43 SAGs (Table S5). Although mitochondrial contigs remained fragmented in most SAGs, four complete or near-complete mitochondrial genomes were recovered with coding content near-identical to the published mitochondrial genome from picozoa MS5584-11^19^ (Fig S9). The ability to assemble complete mitochondrial genomes from the SAGs suggests that the partial nature of the data does not specifically hinder organelle genome recovery if present, at least in the case of mitochondria^20^.

Second, we investigated the possibility that the plastid genome was lost while the organelle itself has been retained—as is the case for *Rhodelphis*^5^. For this, we reconstructed phylogenetic trees for several essential nuclear-encoded biochemical plastid pathways derived by endosymbiotic gene transfer (EGT) that were shown to be at least partially retained even in cryptic plastids^5,21,22^. These included genes involved in the biosynthesis of isoprenoids (ispD,E,F,G,H, dxr, dxs), fatty acids (fabD,F,G,H,I,Z, ACC), heme (hemB,D,E,F,H,Y, ALAS), and iron-sulfur clusters (sufB,C,D,E,S, NifU, iscA; see also Table S6). In all cases, the picozoan homologues grouped either with bacteria—but not cyanobacteria, suggesting contamination—or the mitochondrial/nuclear copies of host origin. Furthermore, none of the picozoan homologues contained predicted N-terminal plastid transit peptides. We also searched for picozoan homologues of all additional proteins (n=62) that were predicted to be targeted to the cryptic plastid in rhodelphids^5^. This search resulted in one protein (Arogenate dehydrogenase, OG0000831) with picozoan homologues that were closely related to red algae and belonged to a larger clade with host-derived plastid targeted plant sequences, but neither the picozoan nor the red algal sequences displayed predicted transit peptides. Finally, to eliminate the possibility of missing sequences because of errors during the assembly and gene prediction, we additionally searched the raw read sequences for the same plastid-targeted or plastid transport machinery genes, which revealed no obvious candidates. In contrast, we readily identified mitochondrial genes (e.g. homologues of the mitochondrial import machinery from the TIM17/TIM22 family), which further strengthened our inference that the single-cell data are in principle adequate to identify organellar components, when they are present.

The lack of cryptic plastids in diverse modern-day picozoans does not preclude photosynthetic ancestry if the plastid was lost early in the evolution of the group. To assess this possibility, we searched more widely for evidence of a cyanobacterial footprint on the nuclear genome that would rise above a background of horizontal gene transfers for proteins functioning in cellular compartments other than the plastids. The presence of a significant number of such proteins may be evidence for a plastid-bearing ancestor. We clustered proteins from 419 genomes, including all major eukaryotic groups as well as a selection of bacteria into orthologous groups (OGs) (Table S7). We built phylogenies for OGs that contained at least cyanobacterial and algal sequences, as well as a sequence from one of 33 focal taxa, including Picozoa, a range of photosynthetic taxa, but also non-photosynthetic plastid-containing, and plastid-lacking taxa to be used as controls. Putative gene transfers from cyanobacteria (EGT) were identified as a group of plastid-bearing eukaryotes that included sequences from the focal taxa and branched sister to a clade of cyanobacteria. We allowed up to 10% of sequences from groups with no plastid ancestry. This approach identified 16 putative EGTs for Picozoa where at least 2 different SAGs/CO-SAGs grouped together, compared to between 89–313 EGTs for photosynthetic species, and up to 59 EGTs for species with non-photosynthetic plastids (Fig 3a). At the other end of the spectrum for species with non-photosynthetic plastids, we observed that the number of inferred cyanobacterial genes for e.g. rhodelphids (14) or *Paraphysomonas* (12) was comparable to Picozoa (16) or other, plastid-lacking taxa such as *Telonema* (15) or *Goniomonas* (18). In order to differentiate these putative endosymbiotic transfers from a background of bacterial transfers (or bacterial contamination), we next attempted to normalise the EGT signal by estimating an extended bacterial signal (indicative of putative HGT: horizontal gene transfers) using the same tree sorting procedure (Fig S10). When comparing the number of inferred EGT with that of inferred HGT, we found a marked difference between plastid-containing (including non-photosynthetic) and plastid-lacking lineages. While all plastid-containing taxa—with the notable exception of *Rhodelphis*—showed a ratio of EGT to HGT above 1, all species without plastid ancestry and *Hematodinium*, one of the few taxa with reported plastid loss, as well as *Rhodelphis* and Picozoa showed a much higher number of inferred HGT than EGT.

**Figure 3.**
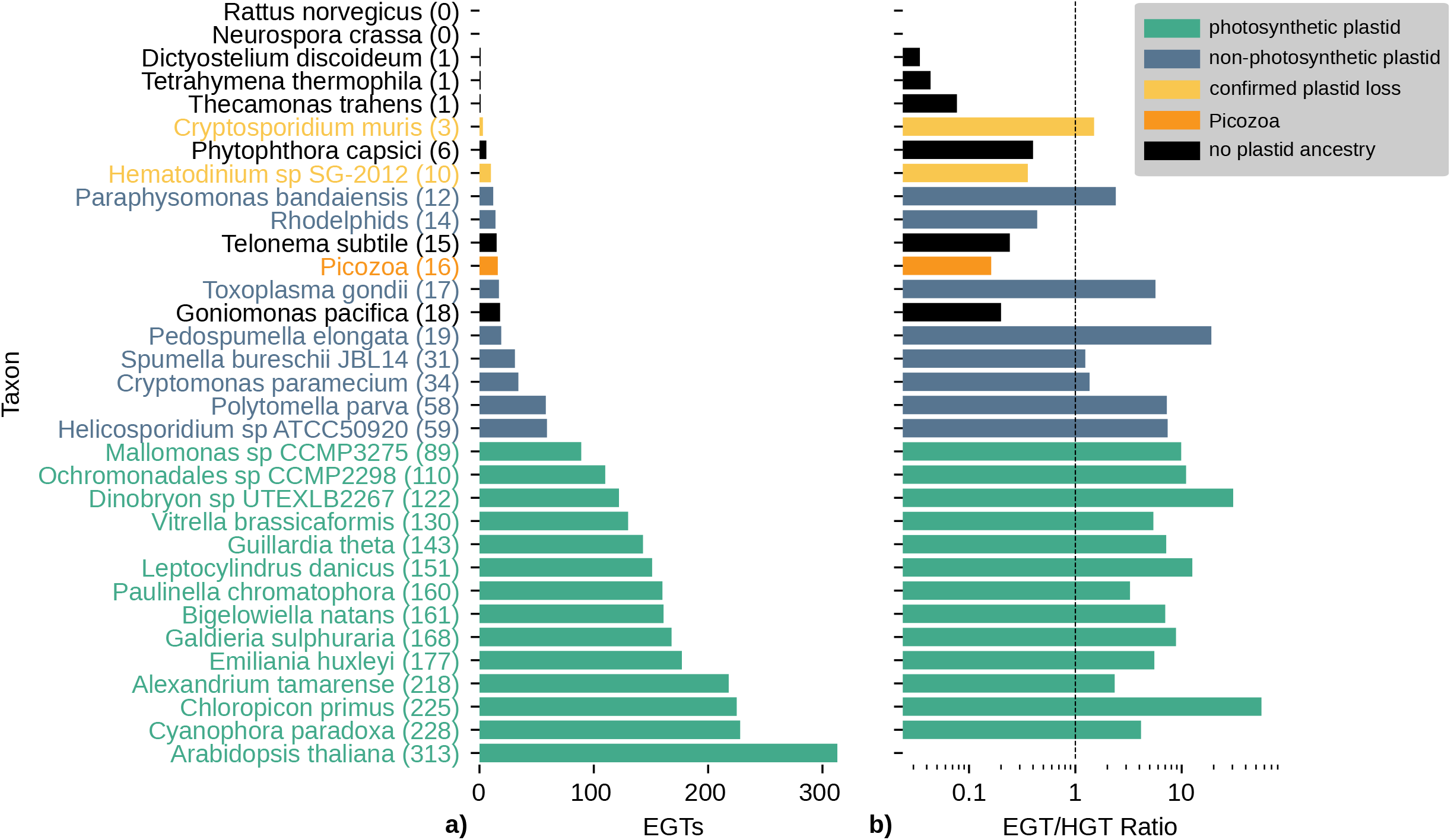
**a)** Number of inferred endosymbiotic gene transfers (EGT) across a selection of 33 species that represent groups with a photosynthetic plastid (green), a non-photosynthetic plastid (blue), confirmed plastid loss (yellow) and no known plastid ancestry (black). These species serve as a comparison to Picozoa (orange). **b)** The number of EGTs from a) is related to the number of inferred HGT across the same 33 selected species. A number below 1 indicates more HGT than EGT, while numbers above 1 indicate more EGT than HGT. No ratio could be calculated for *Arabidopsis* because there were no detectable HGT events.

## Discussion

The 17 SAGs and CO-SAGs of Picozoa obtained in this study provide robust data for phylogenomic analyses of this important phylum of eukaryotes. With this data, we are able to firmly place Picozoa within the supergroup Archaeplastida, most likely as a sister lineage to red algae and rhodelphids. Archaeplastids contain all known lineages with primary plastids (with the exception of *Paulinella*), which are widely viewed to be derived from a single primary endosymbiosis with a cyanobacterium. This notion of a common origin of primary plastids is supported by cellular and genomic data (see^23,24^ and references therein for review), as well as plastid phylogenetics^25,26^. The phylogenetic support for Archaeplastida based on host (nuclear) data has been less certain^7,8,27^, but our analysis is consistent with recent reports that have also recovered a monophyletic origin— here including Picozoa—when using gene and taxon-rich phylogenomic datasets^1,9,10^. This position has important implications for our understanding of plastid origins because, in contrast to all other archaeplastids known to date, our results indicate that Picozoa lack plastids and plastid-associated EGTs. The lack of plastid in Picozoa was also inferred based on smaller initial SAG data^11^ as well as ultrastructural observation of *P. judraskeda*^14^. Two main possible hypotheses exist to explain the lack of plastids in Picozoa: that this group was never photosynthetic, or complete plastid loss occurred early in their evolution.

To suggest that Picozoa was never photosynthetic requires that the current distribution of primary plastids is due to multiple independent endosymbioses, specifically that red algae arose from a separate primary endosymbiosis from that leading to green algae and glaucophytes. This scenario would have involved the endosymbioses of closely related cyanobacterial lineages in closely related hosts to explain the many similarities between primary plastids^24^. Although this may sound unlikely, there is accumulating evidence that similar plastids were derived independently from similar endosymbionts in closely related hosts in dinoflagellates with tertiary plastids^28–30^, and has been argued before for primary plastids^31–34^. However, the current bulk of cell and molecular evidence suggests that multiple independent origins of primary plastids are unlikely, including several features of plastid biology that are not present in cyanobacteria (e.g., protein targeting systems, light-harvesting complex proteins, or plastid genome architecture)^23,24,35^. A related explanation could involve a secondary endosymbiosis where the plastid in red algae, for example, was secondarily acquired from a green alga^36^. This latter scenario would be made unlikely by the identification of host-derived plastid components shared between all archaeplastid lineages.

The second hypothesis implies that a common ancestor of Picozoa entirely lost its primary plastid. The possibility of plastid loss in a free-living lineage like Picozoa would be unprecedented because to date, the only known unambiguous cases of total plastid loss all come from parasitic lineages (all in myzozoan alveolates: in *Cryptosporidium*^37^, certain gregarines^22,38^, and the dinoflagellate *Hematodinium*^39^). To evaluate this possibility, we searched our data for a cyanobacterial footprint in the nuclear genome that would result from an ancestral endosymbiosis. The transfer of genes from endosymbiont to host nucleus via EGT, and the targeting of the product of some or all of these genes back to the plastids, are recognised as a hallmark of organelle integration^40,41^. EGT has occurred in all algae, although its impact on nuclear genomes can vary and the inference of EGT versus other horizontally acquired genes (HGT) can be difficult to decipher for ancient endosymbioses^42–46^. Our analysis of the normalised cyanobacterial signal in Picozoa, which we used as a proxy for quantifying EGT, provides no clear evidence for the existence of a plastid-bearing ancestor. However, it should be noted that evaluating the possibility of plastid loss in groups where a photosynthetic ancestry is not confirmed—such as Picozoa—is complicated because there is no baseline for the surviving footprint of endosymbiosis following plastid loss. Notably, we found no significant difference in the number of inferred EGTs in Picozoa compared to lineages with demonstrated plastid loss (e.g. *Hematodinium* with 10 inferred EGT), lineages with non-photosynthetic plastids (e.g. *Rhodelphis*: 14 inferred EGT), or with no photosynthetic ancestry (e.g. *Telonema*: 15 inferred EGT).

The lack of a genomic baseline to assess plastid loss in Picozoa is further complicated by limitations of our data and methods. The partial nature of eukaryotic SAGs makes it possible that EGTs are absent from our data, even with >90% of inferred genomic completeness. Additionally, the possibility exists that the number of EGT might have always been low during the evolution of the group, even if a plastid was once present. Recent endosymbioses where EGT can be pinpointed with precision showed a relatively low frequency. For example, they represent at most a few percent of the chromatophore proteome in *Paulinella*^47^, or as few as 9 genes in tertiary endosymbiosis in dinoflagellates^48^. Thus, it is possible that the much higher number of EGT inferred in red algae (e.g. 168 in *Galdieria*) occurred after the divergence of Picozoa, and that Picozoa quickly lost its plastid before more EGT occurred. An observation that supports this hypothesis is the low number of putative EGTs found in *Rhodelphis* (14), suggesting that the bulk of endosymbiotic transfers in red algae may have happen after their divergence from rhodelphids.

## Conclusion

In this study, we used single-cell genomics to demonstrate that Picozoa are a plastid lacking major lineage of archaeplastids. This is the first example of an archaeplastid lineage without plastids, which can be interpreted as either plastid loss, or evidence of independent endosymbiosis in the ancestor of red algae and rhodelphids. Under the most widely accepted scenario of a single plastid origin in Archaeplastida, Picozoa would represent the first known case of plastid loss in this group, but also more generally in any free-living species. In order to discriminate plastid loss from multiple plastid gains in the early archaeplastid evolution, and more generally during the evolution of secondary or tertiary plastids, a better understanding of the early steps of plastid integration is required. In the recently evolved primary plastid-like chromatophores of *Paulinella*, the transfer of endosymbiotic genes at the onset of the integration was shown to be minimal^4^. Similar examples of integrated plastid endosymbionts but with apparently very few EGTs are known in dinoflagellates^48,49^. Therefore, new important clues to decipher the origin of plastids will likely come from a better understanding of the role of the host in driving these endosymbioses, and crucially the establishment of a more complete framework for archaeplastid evolution with the search and characterization of novel diversity of lineages without plastids. The fact that this lineage has never been successfully maintained in culture, with just one study achieving transient culture^14^, might indicate a lifestyle involving close association with other organisms (such as symbiosis) and further underscores the enigma of picozoan biology, the lack of information on which hinders our interpretation of their evolution.

## Material and Methods

### Cell Isolation, Identification, and genome amplification

#### Baltic Sea

Surface (depth: up to 2 m) marine water was collected from the Linnaeus microbial Observatory (LMO) in the Baltic Sea located at 56°N 55.85’ and 17°E 03.64’ on two occasions: 2 May 2018 (6.1°C and 6.8 ppt salinity) and 3 April 2018 (2.4°C and 6.7 ppt salinity). The samples were transported to the laboratory and filter-fractionized (see Table 1 for detailed sample information). The size fractions larger than 2 µm were discarded whereas the fraction collected on 0.2 µm filters was resuspended in 2 mL of the filtrate. The obtained samples were used for fluorescence-activated cell sorting (FACS). Aliquots of 4 µL of 1 mM Mitotracker Green FM (ThermoFisher) stock solution were added to the samples and were kept in the dark at 15°C for 15–20 minutes. Then the cells were sorted into empty 96-well plates using MoFlo Astrios EQ cell sorter (Beckman Coulter). Gates were set mainly based on Mitotracker intensity and the dye was detected by a 488 nm and 640 nm laser for excitation, 100 µm nozzle, sheath pressure of 25 psi and 0.1 µm sterile filtered 1 x PBS as sheath fluid. The region with the highest green fluorescence and forward scatter contained the target group and was thereafter used alongside with exclusion of red autofluorescence.

The SAGs were generated in each well with REPLI-g® Single Cell kit (Qiagen) following the manufacturer’s recommendations but scaled down to 5 µl reactions. Since the cells were sorted in dry plates, 400 nl of 1xPBS was added prior to 300 nl of lysis buffer D2 for 10 min at 65°C and 10 min on ice, followed by 300 nl stop solution. The PBS, reagent D2, stop solution, water, and reagent tubes were UV-treated at 2 Joules before use. A final concentration of 0.5 µM SYTO 13 (Invitrogen) was added to the MDA mastermix. The reaction was run at 30°C for 6 h followed by inactivation at 65°C for 5 min and was monitored by detection of SYTO13 fluorescence every 15 minutes using a FLUOstar® Omega plate reader (BMG Labtech, Germany). The single amplified genome (SAG) DNA was stored at -20°C until further PCR screening. The obtained products were PCR-screened using Pico-PCR approach, as described in^14^ and the wells showing signal for Picozoa were selected for sequencing.

#### Eastern North Pacific

Seawater was collected and sorted using a BD InFlux Fluorescently Activated Cell Sorter (FACS) on three independent cruises in the eastern North Pacific. The instrument was equipped with a 100 mW 488 nm laser and a 100 mW 355 nm laser and run using sterile nuclease-free 1× PBS as sheath fluid. The stations where sorting occurred were located at 36.748°N, 122.013°W (Station M1; 20 m, 2 April 2014 and 10 m, 5 May 2014); 36.695°N, 122.357°W (Station M2, 10 m, 5 May 2014); and 36.126°N, 123.49°W (Station 67-70, 20 m 15 October 2013). Water was collected using Niskin bottles mounted on a CTD rosette. Prior to sorting samples were concentrated by gravity over a 0.8 μm Supor filter. Two different stains were used: LysoSensor (2 April 2014, M1) and LysoTracker (5 May 2014, M1; 15 October 2013, 67-70), or both together (5 May 2014, M2). Selection of eukaryotic cells stained with LysoTracker Green DND-26 (Life Technologies; final concentration, 25 nM) was based on scatter parameters, positive green fluorescence (520/35 nm bandpass), as compared to unstained samples, and exclusion of known phytoplankton populations, as discriminated by their forward angle light scatter and red (chlorophyll-derived) autofluorescence (i.e., 692/40 nm bandpass) under 488 nm excitation, similar to methods in^50^. Likewise, selection of cells stained with LysoSensor Blue DND-167 (Life Technologies; final concentration, 1 μM), a ratiometric probe sensitive to intracellular *pH* levels, e.g. in lysosomes, was based on scatter parameters, positive blue fluorescence (435/40 nm bandpass), as compared to unstained samples, and exclusion of known phytoplankton populations, as discriminated by their forward angle light scatter and red (chlorophyll-derived) autofluorescence (i.e., 692/40 nm bandpass filter) under 355 nm excitation. For sorts using both stains, all of the above criteria, and excitation with both lasers (with emissions collected through different pinholes and filter sets), were applied to select cells. Before each sort was initiated, the respective plate was illuminated with UV irradiation for 2 min. Cells were sorted into 96- or 384-well plates using the Single-Cell sorting mode from the BD FACS Software v1.0.0.650. A subset of wells was left empty or received 20 cells for negative and positive controls, respectively. After sorting, the plates were covered with sterile, nuclease free foil and frozen at -80 °C immediately after completion.

Whole genome amplification of individual sorted cells followed methods outlined in^50^. For initial screening, 18S rRNA gene amplicons were amplified from each well using the Illumina adapted TAReuk454FWD1 and TAReukREV3 primers targeting the V4 hypervariable region. PCR reactions contained 10 ng of template DNA and 1X 5PRIME HotMasterMix (Quanta Biosciences) as well as 0.4 mg ml^-1^ BSA (NEB) and 0.4 μM of each primer. PCR reactions entailed: 94 °C for 3 min; and 30 cycles at 94 °C for 45 sec, 50 °C for 60 sec and 72 °C for 90 sec; with a final extension at 72 °C for 10 min. Triplicate reactions per cell were pooled prior to Paired-end (PE) library sequencing (2 × 300 bp) and the resulting 18S V4 rRNA gene amplicons were trimmed at Phred quality (Q) of 25 using a 10 bp running window using Sickle 1.33 (https://github.com/najoshi/sickle). Paired-end reads were merged using USEARCH v.9.0.2132 when reads had a ≥40 bp overlap with max 5% mismatch. Merged reads were filtered to remove reads with maximum error rate >0.001 or <200 bp length. Sequences with exact match to both primers were retained, primer sequences were trimmed using Cutadapt v.1.13^51^, and the remaining sequences were *de novo* clustered at 99% sequence similarity by UCLUST forming operational taxonomic units (OTUs). Each of the cells further sequenced had a single abundant OTU that was taxonomically identified using BLASTn in GenBank’s nr database.

### Sequencing

Sequencing libraries were prepared from 100 ng DNA using the TruSeq Nano DNA sample preparation kit (cat# 20015964/5, Illumina Inc.) targeting an insert size of 350bp. For six samples, less than 100ng was used (between 87 ng-97 ng). The library preparations were performed by SNP&SEQ Technology Platform at Uppsala University according to the manufacturers’ instructions. All samples were then multiplexed on one lane of an Illumina HiSeqX instrument with 150 cycles paired-end sequencing using the v2.5 sequencing chemistry, producing between 10,000 and 30,000,000 read pairs.

### Genome Assembly and 18S rRNA gene analysis

The 43 Illumina datasets were trimmed using *Trim Galore* v0.6.1 (https://www.bioinformatics.babraham.ac.uk/projects/trim_galore/) with default parameter and assembled into genomic contigs with SPAdes v3.13.0^52^ in single-cell mode (--sc --careful -k 21,33,55,99). Open reading frames (ORFs) were identified and translated using Prodigal v2.6.3 in ‘anonymous’ mode^53^ and rRNA genes were predicted using barrnap v0.9 (https://github.com/tseemann/barrnap) for eukaryotes. All 18S rRNA gene sequences were, together with available reference sequences from the protist ribosomal reference database (PR2), aligned with MAFFT E-INS-i v7.429^54^ and trimmed with trimal^55^ (gap threshold 0.01%). After performing a modeltest using ModelFinder^56^ (best model: GTR+R6+F), a phylogenetic tree was reconstructed in IQ-TREE v2.1.1^57^ with 1000 ultrafast bootstrap replicates (see Fig S11 for a tree with extended taxon sampling). Additionally, we estimated the average nucleotide identity (ANI) for all pairs of SAGs using fastANI v1.2^58^ (Fig S12). Based on the 18S rRNA gene tree and the ANI value, groups of closely related SAGs with almost identical 18S rRNA gene sequences (sequence similarity above 99%) were identified for co-assembly. Co-assemblies were generated in the same way as described above for single assemblies, pooling sequencing libraries from closely related single cells. ORFs and rRNA genes were similarly extracted from the co-assemblies. The completeness of the SAGs and CO-SAGs was then assessed using BUSCO v4.1.3^15^ with 255 markers for eukaryotes (Fig S13) as well as using the 320 marker phylogenomic dataset as described below. General genome characteristics were computed with QUAST v5.0.2^59^. Alignments were reconstructed for the 18S rRNA genes from the co-assemblies and those SAGs not included in any CO-SAG together with PR2 references for cryptists and katablepharids (the closest groups to Picozoa in 18S rRNA gene phylogenies) in the same way as described above. The tree was reconstructed using GTR+R4+F after model selection and support was assessed with 100 non-parametric bootstraps. The six CO-SAGs and the 11 individual SAGs were used in all subsequent analyses.

For each of these 17 assemblies we estimated the amount of prokaryotic/viral contamination by comparing the predicted proteins against the NCBI nr database using DIAMOND in blastp mode^60^. If at least 60% of all proteins from a contig produced significant hits only to sequences annotated as prokaryotic or viral, we considered that contig to be a putative contamination. In general only a small fraction of each assembly was found to be such a contamination (Fig S14).

### Phylogenomics

Existing untrimmed alignments for 320 genes and 763 taxa from^1^ were used to create HMM profiles in HMMER v3.2.1^61^, which were then used to identify homologous sequences in the protein sequences predicted from the Picozoa assemblies (or co-assemblies) as well as in 20 additional, recently sequenced eukaryotic genomes and transcriptomes (Table S2). Each single gene dataset was filtered using PREQUAL v1.02^62^ to remove non-homologous residues prior to alignment, aligned using MAFFT E-INS-i, and filtered with Divvier -partial v1.0^63^. Alignments were then used to reconstruct gene trees with IQ-TREE (-mset LG, LG4X; 1000 ultrafast bootstraps with the BNNI optimization). All trees were manually scrutinized to identify contamination and paralogs. These steps were repeated at least two times, until no further contaminations or paralogs could be detected. We excluded three genes that showed ambiguous groupings of Picozoa or rhodelphids in different parts of the trees. From this full dataset of 317 genes and 794 taxa, we created a concatenated supermatrix alignment using the cleaned alignments described above. This supermatrix was used to reconstruct a tree in IQ-TREE with the model LG+G+F and ultrafast bootstraps (1000 UFBoots) estimation with the BNNI improvement.

We then prepared a reduced dataset with a more focused taxon sampling of 67 taxa, covering all major eukaryotic lineages but focussing on the groups for which an affiliation to Picozoa had been reported previously. For this dataset, closely related species were merged into OTUs in some cases in order to decrease the amount of missing data per taxon (Table S3). The 317 single gene datasets were re-aligned using MAFFT E-INS-i, filtered using both Divvier -partial and BMGE (-g 0.2 -b 10 -m BLOSUM75) and concatenated into two supermatrices. Model selection of mixture models was performed using ModelFinder^56^ for both datasets, and in both cases LG+C60+G+F was selected as the best-fitting model. Trees for both datasets were reconstructed using the Posterior Mean Site Frequency (PMSF)^64^ approximation of this mixture model in IQ-TREE and support was assessed with 100 non-parametric bootstraps (see Fig S15 for the Divvier derived tree)

In addition, we reconstructed a phylogenetic tree using the supermatrix alignment based on BMGE trimming in PhyloBayes MPI v1.8^65^ using the CAT+GTR+G model. We ran three independent chains for 3600 cycles, with the initial 1500 cycles being removed as burnin from each chain. We then generated a consensus tree using the bpcomp program of PhyloBayes. Partial convergence was achieved between chains 1 and 2 with a maxdiff value of 0.26 (Fig S16). The third chain differed only in the position of haptists and *Ancoracysta twista*, but not in the relationships within Archaeplastida and the position of Picozoa (Fig S17).

In order to test the robustness of our results we additionally performed a fast-site removal analysis^66^, iteratively removing the 5000 fastest evolving sites (up to a total of 55000 removed sites). For each of these 11 alignments, we reconstructed an ML tree using the model LG+C60+G+F in IQ-TREE with ultrafast bootstraps (1000 UFBoots) and evaluated the support for the branching of Picozoa with rhodelphids and red algae as well as for other groupings (Fig S4). We also performed trimming of the 25% and 50% most heterogeneous sites based on the χ2 metric^67^ and performed tree reconstruction using the same model as above (Fig S5, Fig S7). We also prepared a supermatrix alignment (BMGE trimmed) from 224 genes with at least two Picozoa sequences in the final dataset and performed similar tree reconstruction (model LG+C60+G+F in IQ-TREE with 1000 ultrafast bootstraps, Fig S8).

Furthermore, we performed a supertree-based phylogenetic reconstruction using ASTRAL-III v5.7.3^*68*^. We reconstructed gene trees for each of the 317 alignments of the 67-taxa dataset using IQ-TREE (-m TEST –mset LG -mrate G,R4 -madd LG4X,LG4X+F,LG4M,LG4M+F, using 1000 ultrafast bootstraps) and performed multi-locus bootstrapping based on the bootstrap replicates (option -b in ASTRAL-III) (Fig S6).

Finally, we performed an approximately unbiased (AU) test in IQ-TREE of 15 topologies (see Table S4), including previously recovered positions of Picozoa (as sister to red algae, cryptists, telonemids, Archaeplastida etc.).

### Mitochondrial contig identification and annotation

Using the published picozoan mitochondrial genome (Picozoa sp. MS584-11: MG202007.1 from^19^), BLAST searches were performed on a dedicated sequenceServer^69^ to identify mitochondrial contigs in the 43 picozoan SAGs. Putative mitochondrial contigs were annotated using the MFannot server (https://megasun.bch.umontreal.ca/cgi-bin/mfannot/mfannotInterface.pl). All contigs with predicted mitochondrial genes or whose top hits in the NCBI nr database was the published picozoan mitochondrial genome (MG202007.1) were considered to be *bona fide* mitochondrial contigs and retained (Supplementary materials). Manual annotation was conducted as needed.

### Plastid Genes & EGT

GetOrganelle v1.7.1^70^ was used to identify organellar genomes. We searched the assemblies for putative plastid contigs with the subcommand ‘get_organelle_from_assembly.py -F embplant_pt,other_pt’, while we attempted to assemble such a genome directly using the command ‘get_organelle_from_reads.py -R 30 -k 21,45,65,85,105 -F embplant_pt,other_pt’. We additionally searched the predicted proteins against available plastid protein sequences from ncbi using DIAMOND v2.0.6^60^ in blastp mode (--more-sensitive). Contigs that were identified as putatively coming from a plastid genome were then checked manually by doing BLAST searches against NT, and contigs that showed similarity only to bacterial genomes or to the picozoa mitochondrial assembly MG202007.1 were rejected.

To search for known plastid pathways, we prepared Hidden Markov model (HMM) profiles for 32 gene alignments that were shown to be retained in lineages with non-photosynthetic plastids and included a wide diversity of plastid-bearing eukaryotes following a similar approach as in^22^. Using these profiles, we identified homologues in the Picozoa SAGs, and aligned them together with the initial sequences used to create the profiles using MAFFT E-INS-i. We trimmed the alignments using trimAl v1.4.rev15 ‘-gt 0.05’ and reconstructed phylogenetic trees using IQ-TREE (-m LG4X; 1000 ultrafast bootstraps with the BNNI optimization) from these alignments. We then manually inspected the trees to assess whether picozoan sequences grouped with known plastid-bearing lineages. We additionally used the sequences from these core plastid genes to search the raw sequencing reads for any signs of homologues that could have been missed in the assemblies. We used the tool PhyloMagnet^71^ to recruit reads and perform gene-centric assembly of these genes^72^. The assembled genes were then compared to the NR database using DIAMOND in blastp mode (--more-sensitive --top 10).

To identify putative EGT, we prepared orthologous clusters for 419 species (128 bacteria and 291 eukaryotes) with a focus on plastid-bearing eukaryotes and cyanobacteria, but also including other eukaryotes and bacteria, using OrthoFinder v2.4.0^73^. For Picozoa and a selection of 32 photosynthetic or heterotrophic lineages (Table S8), we inferred trees for 2626 clusters that contained the species under consideration, at least one cyanobacterial sequence, and at least one archaeplastid sequence of red algae, green algae or plants. Alignments for these clusters were generated with MAFFT E-INS-i, filtered using trimAl ‘-gt 0.01’ and phylogenetic trees were reconstructed using IQ-TREE (-m LG4X; 1000 ultrafast bootstraps with the BNNI optimization). We then identified trees where the target species grouped with other plastid-bearing lineages (allowing up to 10% non-plastid sequences) and sister at least two cyanobacterial sequences. For Picozoa, we added the condition that sequences from at least two SAG/COSAG assemblies must be monophyletic. For species with no known plastid ancestry such as *Rattus* or *Phytophthora*, putative EGTs can be interpreted as false positives due to contamination, poor tree resolution or other mechanisms, since we expect no EGTs from cyanobacteria to be present at all in these species. This rough estimate of the expected false positive rate for this approach can give us a baseline of false positives that can be expected for picozoa as well.

To put the number of putative EGTs into relation to the overall amount of gene transfers, we applied a very similar approach to the one described above for detecting putative HGT events. We prepared additional trees (in the same way as described for the detection of EGTs) for clusters that contained the taxon of interest and non-cyanobacterial bacteria and identified clades of the taxon under consideration (including a larger taxonomic group, e.g. Streptophyta for *Arabidopsis* or Metazoa for *Rattus*) that branched sister to a bacterial clade.

### *Distribution of Picozoa in* Tara *Oceans*

We screened available OTUs that were obtained from V9 18S rRNA gene eukaryotic amplicon data generated by *Tara* Oceans^17^ for sequences related to Picozoa. Using the V9 region of the 18S rRNA gene sequences from the 17 Picozoa assemblies as well as from the picozoan PR2 references used to reconstruct the 18S rRNA gene tree described above, we applied VSEARCH v2.15.1^74^ (--usearch_global -iddef 1 --id 0.90) to find all OTUs with at least 90 % similar V9 regions to any of these reference picozoan sequences. Using the relative abundance information available for each *Tara* Oceans sampling location, we then computed the sum for all identified Picozoa OTUs per station and plotted the relative abundance on a world map.

## Acknowledgments

This work was supported by a grant from Science for Life Laboratory available to FB and a scholarship from Carl Tryggers Stiftelse to VZ (PI: FB). TJGE thanks the European Research Council (ERC consolidator grant 817834); the Dutch Research Council (NWO-VICI grant VI.C.192.016); Moore–Simons Project on the Origin of the Eukaryotic Cell (Simons Foundation 735925LPI, https://doi.org/10.46714/735925LPI); and the Marie Sklodowska-Curie ITN project SINGEK (H2020-MSCA-ITN-2015-675752) which provided funding for MES. PJK and VM were funded by an Investigator Grant from the Gordon and Betty Moore Foundation (https://doi.org/10.37807/GBMF9201). The Pacific Ocean work was supported by GBMF3788 to AZW. Sampling at the LMO station in the Baltic Sea was carried out by support from the Swedish Research Council VR and the marine strategic research program EcoChange to JP. Sequencing was performed by the SNP&SEQ Technology Platform in Uppsala, part of the National Genomics Infrastructure (NGI) Sweden and Science for Life Laboratory. The SNP&SEQ Platform is also supported by the Swedish Research Council and the Knut and Alice Wallenberg Foundation. Cell sorting and whole genome amplification was performed at the Microbial Single Cell Genomics Facility (MSCG) at SciLifeLab. Computations were performed on resources provided by the Swedish National Infrastructure for Computing (SNIC) at Uppsala Multidisciplinary Center for Advanced Computational Science (UPPMAX) under Projects SNIC 2019/3-305, SNIC 2020/15-58, SNIC 2021/5-50, Uppstore2018069. Finally, we thank Eunsoo Kim and Sally D. Warring for sharing peptide models from *Palpitomonas bilix* and *Roombia truncata*.

## Author contributions

FB and JGW conceived the study. For the Baltic samples, VVZ and JP performed sampling and cell sorting; VVZ and JFHS performed genome amplification & sequencing preparation. For the Pacific samples, CP, SW and AZW conceived, developed and implemented sort protocols; performed sampling, cell sorting and sequencing; CP and AZW performed initial sequence analyses and phylogenetics. MES under the supervision of TJGE and FB performed assembly of SAGs and Co-SAGs, phylogenomic analyses, and searched for plastid evidence and gene transfers with the help of VM and PJK. RPS and JGW assembled mitochondrial genomes. MES, FB, and JGW drafted the manuscript. TJGE, PKJ, AZW, JP, VVZ and JFHS contributed edits to the manuscript. All authors read and approved the final version.

## Code & data availability

All custom scripts used in this study are available at https://github.com/maxemil/picozoa-scripts under a MIT license. All data used for the analyses as well as results files such as contigs and single gene trees are available at figshare (10.6084/m9.figshare.c.5388176). A sequenceServer BLAST server was set up for the SAG assemblies: http://evocellbio.com/SAGdb/burki/. Raw sequencing reads were deposited in the Sequence Read Archive (SRA) at NCBI under accession PRJNA747736 and will be available upon acceptance.

