## Supplementary Figures for "Single cell genomics reveals plastid-lacking Picozoa are close relatives of red algae"

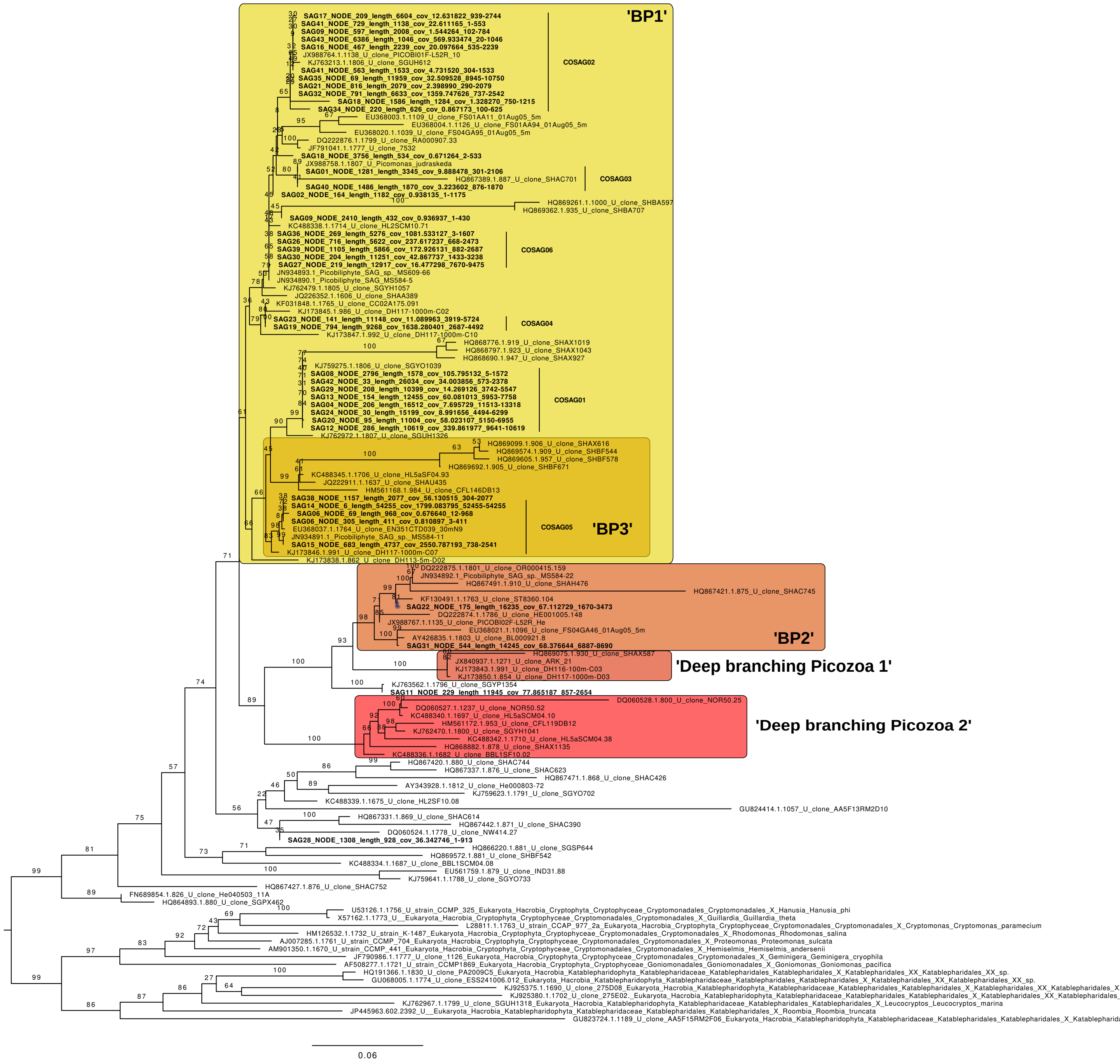

**Supplementary Figure 1: Maximum Likelihood tree of the 18S rRNA gene.** Sequences from the individual SAG assemblies and a number of reference sequences from Picozoa, cryptophyes katablepharids from the PR2 database. The tree was reconstructed under the model GTR+G.

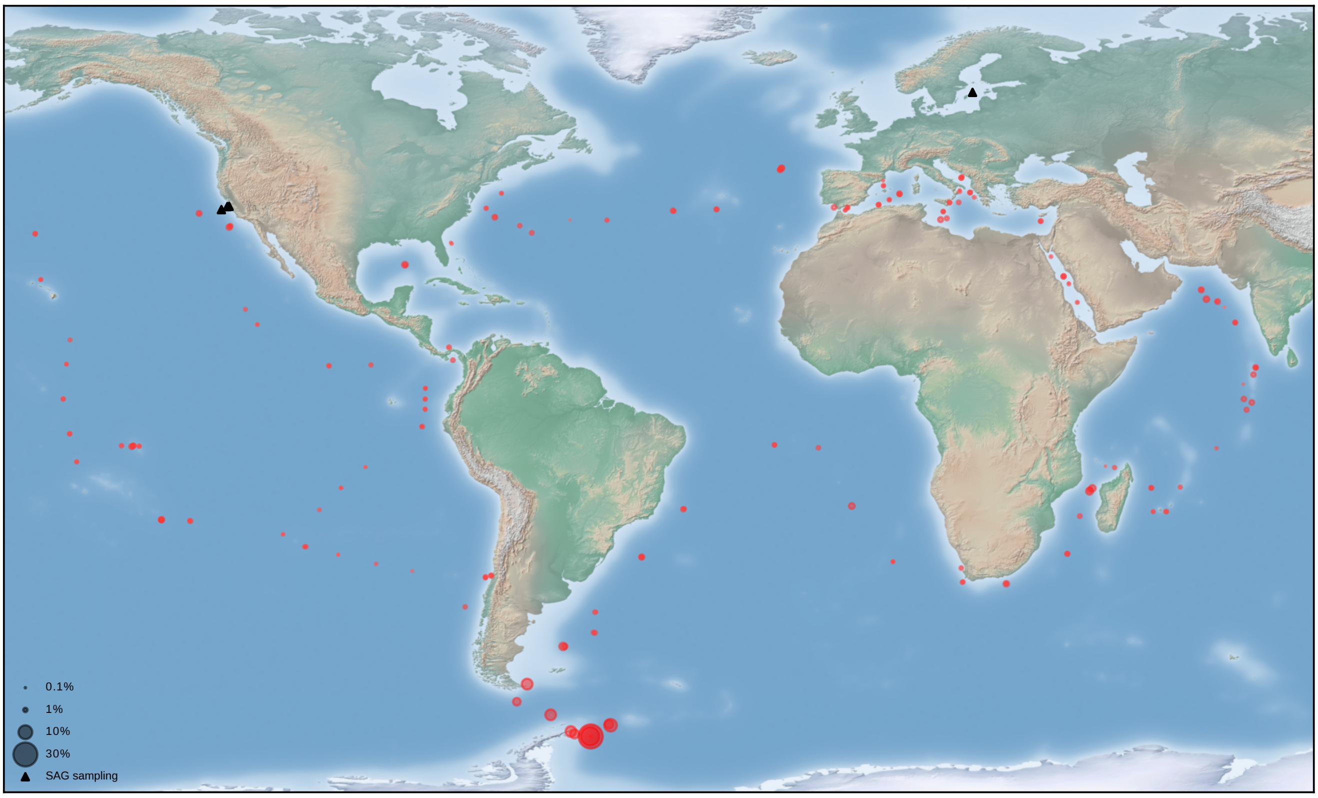

**Supplementary Figure 2: Combined relative abundance of all Picozoa OTUs identified in the Tara Oceans metabarcoding data.** Abundances are given on a corresponding map of sampling locations. Size of the circles corresponds to relative abundance. Triangles mark the location of single-cell sampling from this study.

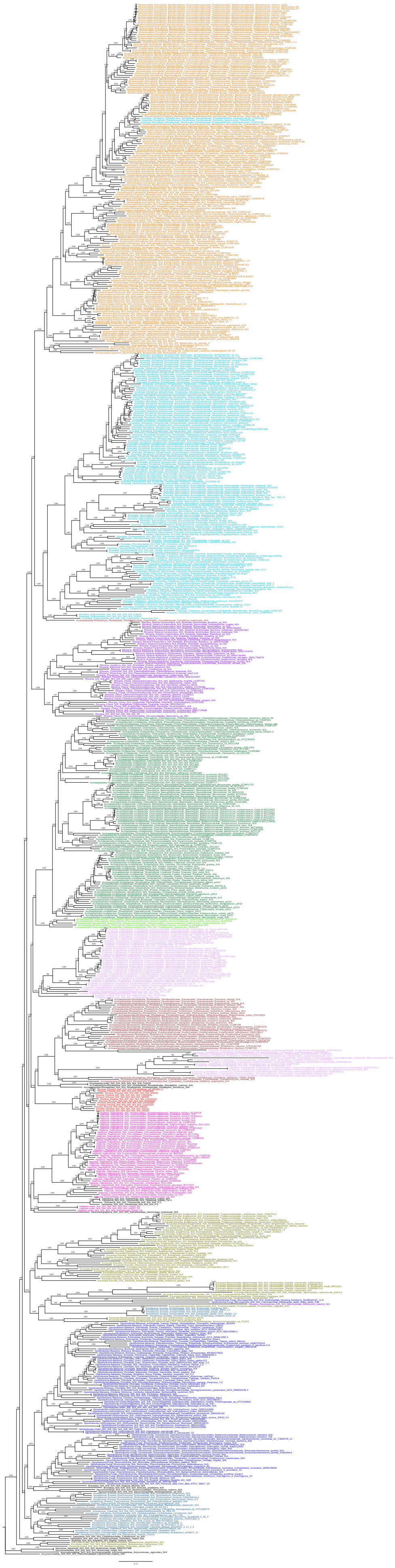

**Supplementary Figure 3: Maximum likelihood tree of 794 eukaryotic species.** The tree is based on the concatenated alignment of 317 marker genes (filtered with Divvier) and was reconstructed using the site-homogeneous model LG+F+G. Support values correspond to 1000 ultrafast bootstrap replicates.

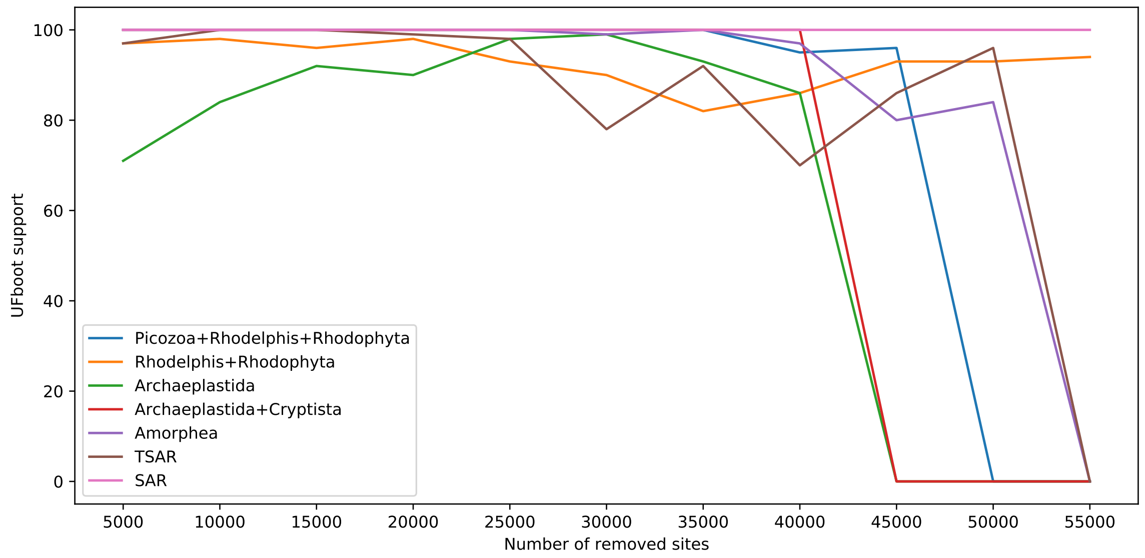

**Supplementary Figure 4: Support for several groupings as estimated in different trees with increasing number of fast-evolving sites removed.** Initial alignment was the 67-dataset (filtered with BMGE). All trees were reconstructed using the site-heterogeneous model LG+C60+F+G, support values correspond to 1000 ultrafast bootstrap replicates

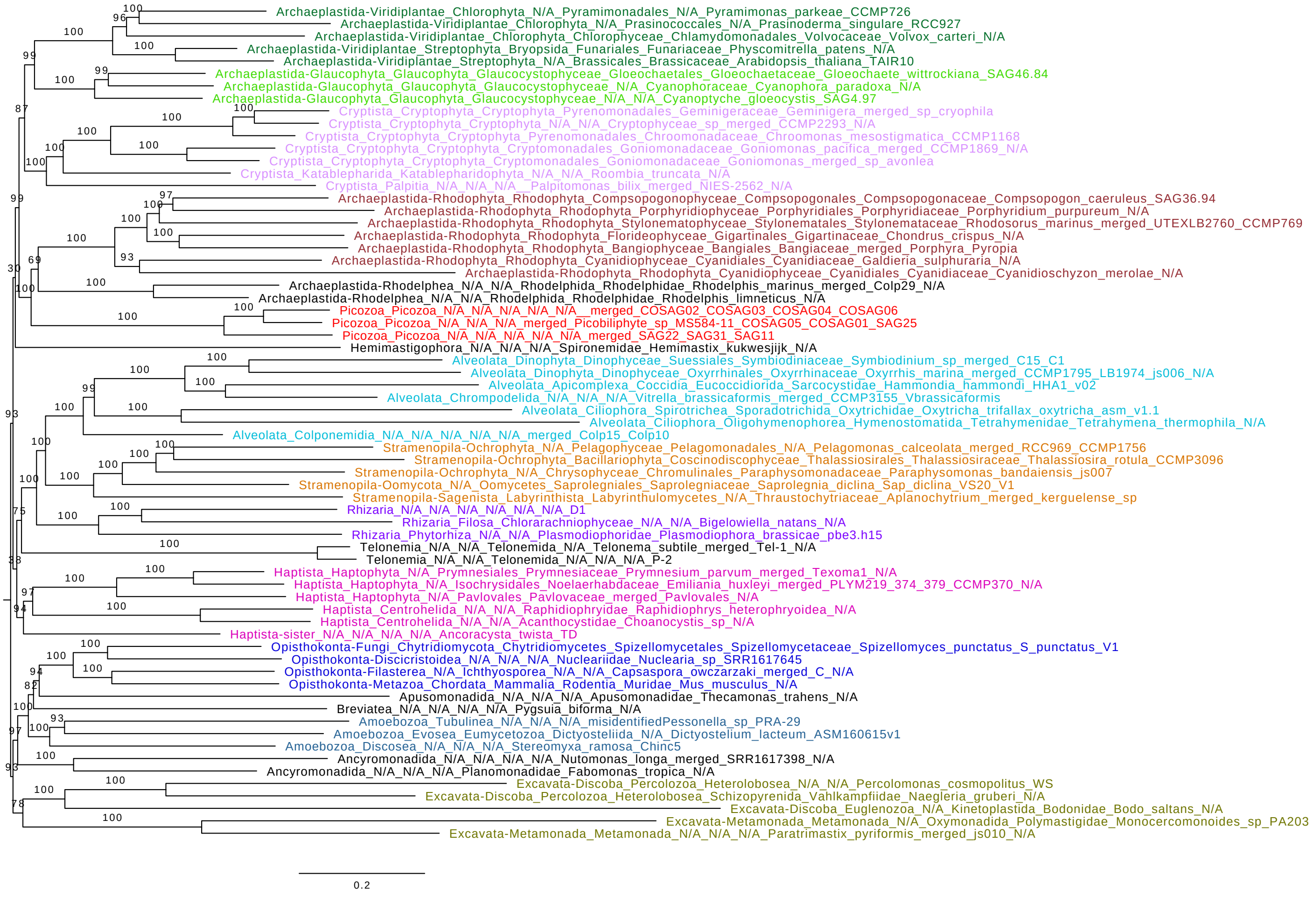

**Supplementary Figure 5: Maximum likelihood tree of 67 eukaryotic species showing the position of Picozoa.** The tree is based on the concatenated alignment of 317 marker genes, filtered with BMGE and trimmed of the 25% most heterogenous sites according to the chi-square statistic. The tree was reconstructed using the site-heterogeneous model LG+C60+F+G, support values correspond to 1000 ultrafast bootstrap replicates.

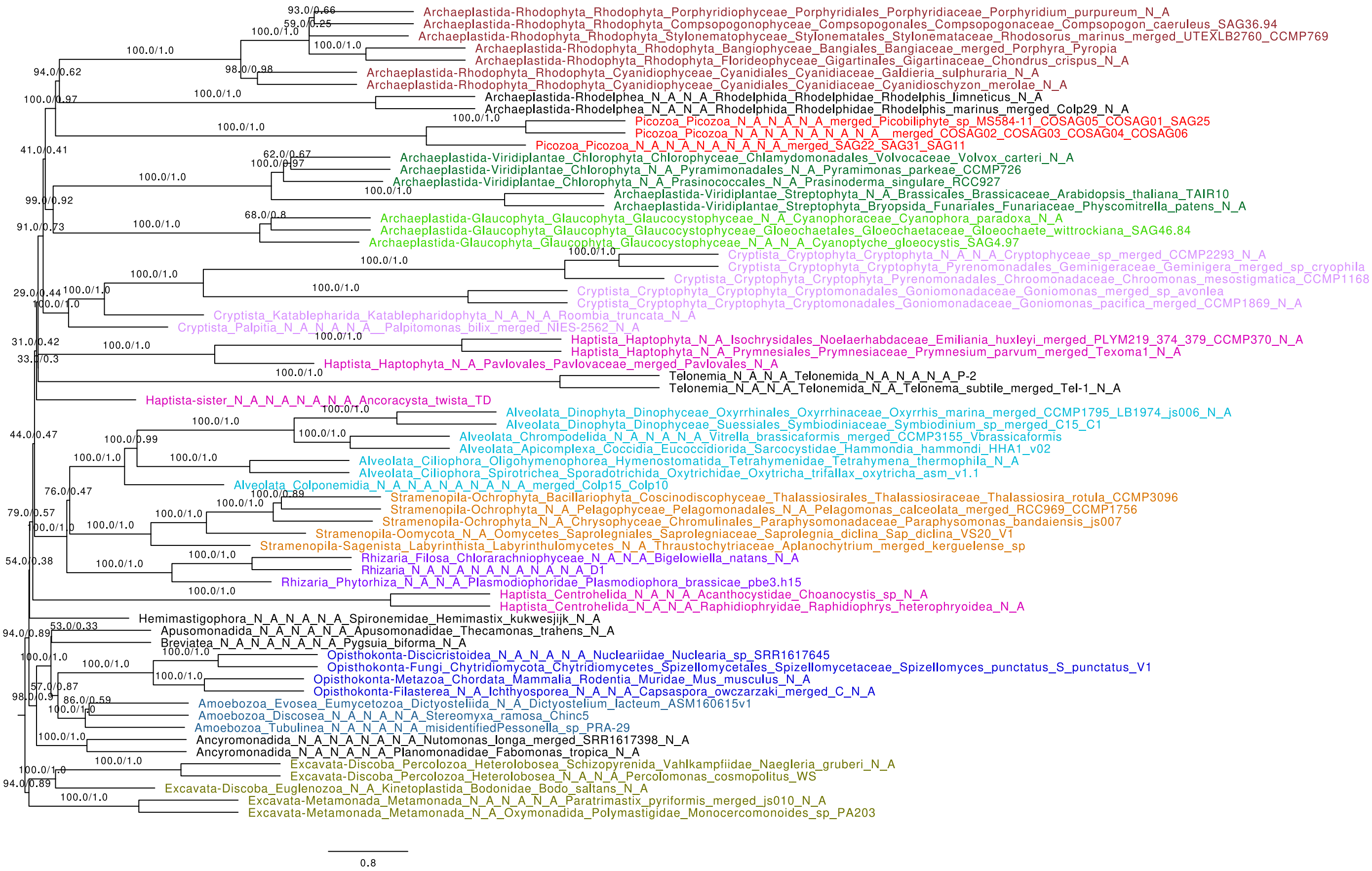

**Supplementary Figure 6: Multi-species coalescent species tree reconstructed with ASTRAL-III.** Individual single gene trees were inferred for all 317 untrimmed alignments from the 67-taxa dataset, using automatic model selection and 1000 ultrafast bootstraps. Support values correspond to multi-locus bootstraps and posterior probabilities, respectively.

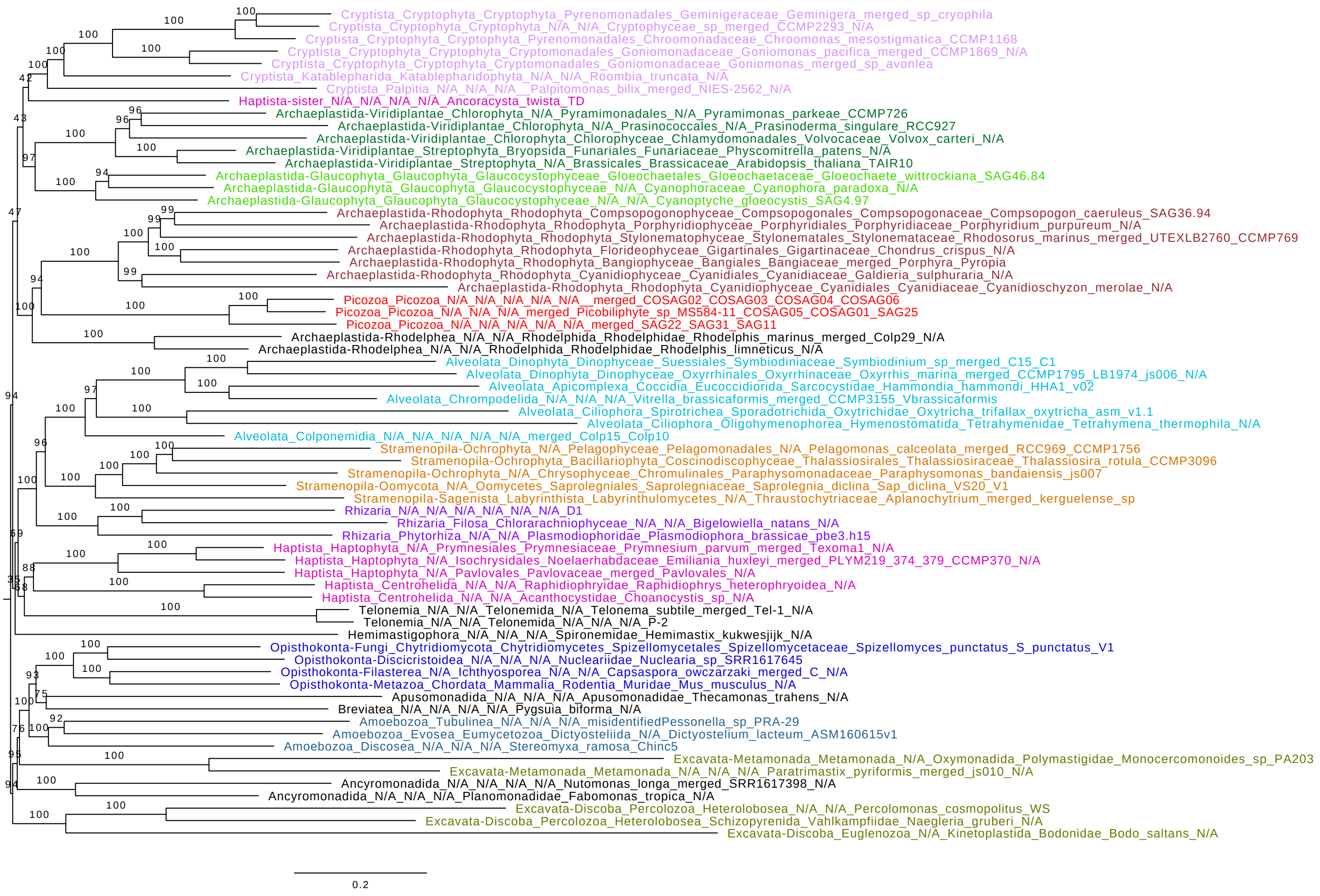

**Supplementary Figure 7: Maximum likelihood tree of 67 eukaryotic species showing the position of Picozoa.** The tree is based on the concatenated alignment of 317 marker genes, filtered with BMGE and trimmed of the 50% most heterogenous sites according to the chi-square statistic. The tree was reconstructed using the site-heterogeneous model LG+C60+F+G, support values correspond to 1000 ultrafast bootstrap replicates.

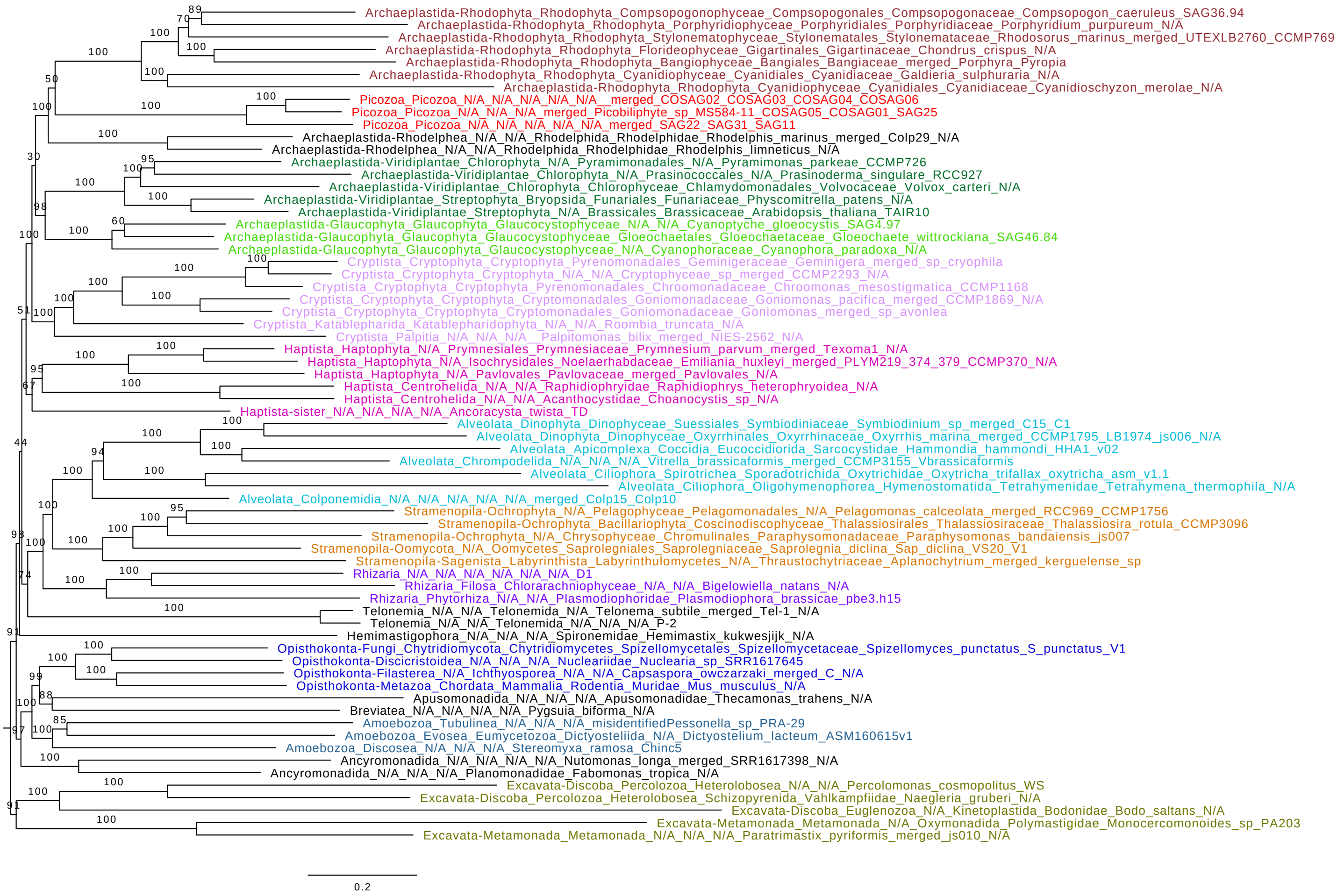

**Supplementary Figure 8: Maximum likelihood tree of 67 eukaryotic species showing the position of Picozoa.** The tree is based on the concatenated alignment of 224 marker genes with at least two monophyletic Picozoa sequences (filtered with BMGE). The tree was reconstructed using the site-heterogeneous model LG+C60+F+G, support values correspond to 1000 ultrafast bootstrap replicates.

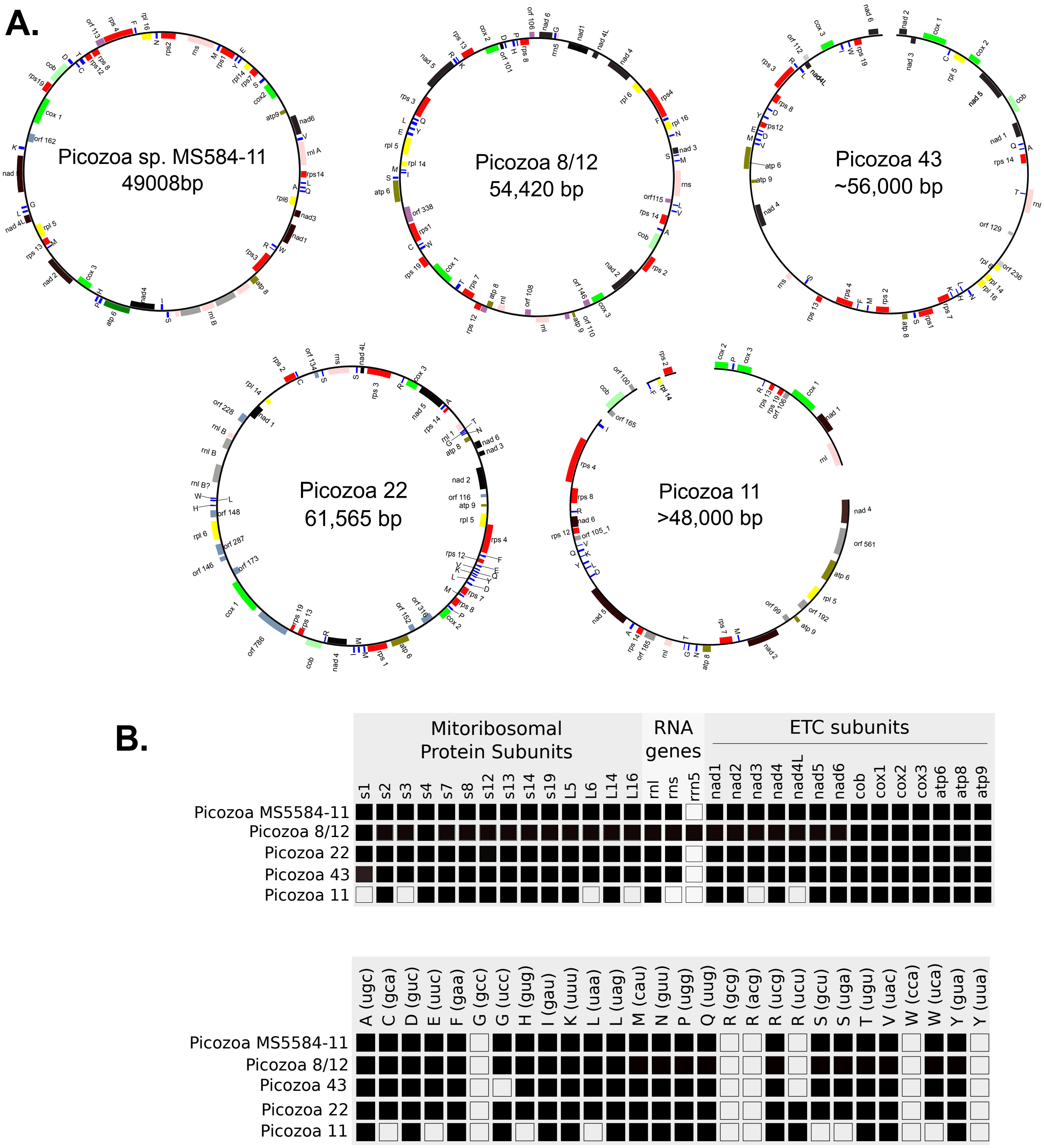

**Supplementary Figure 9: Complete and near complete mitochondrial genomes assembled from diverse picozoan SAGs.** A. Circular depictions of picozoan mitochondrial genomes compared to the published Picozoa MS584-11 sequence (Janouskovec et al. 2017). Mitochondrial contigs were annotated using mfannot with manual corrections as needed (http://megasun.bch.umontreal.ca/RNAweasel/). MtDNAs are represented as circular diagrams or broken circles if contigs could not be joined. For SAGs 8 and 12, the mitochondrial genome was inferred by stitching near-identical stretches together. Colour-coded genes: black, complex I; green, complex III; light green, complex IV; dark green, complex V; yellow, small ribosomal subunit proteins; red, large ribosomal subunit proteins; blue, tRNAs; pink, rnl and rns genes. B. gene complement of sequenced picozoan mitochondrial genomes. Top: protein coding and ribosomal RNA gene complement. Bottom: tRNA gene complement.

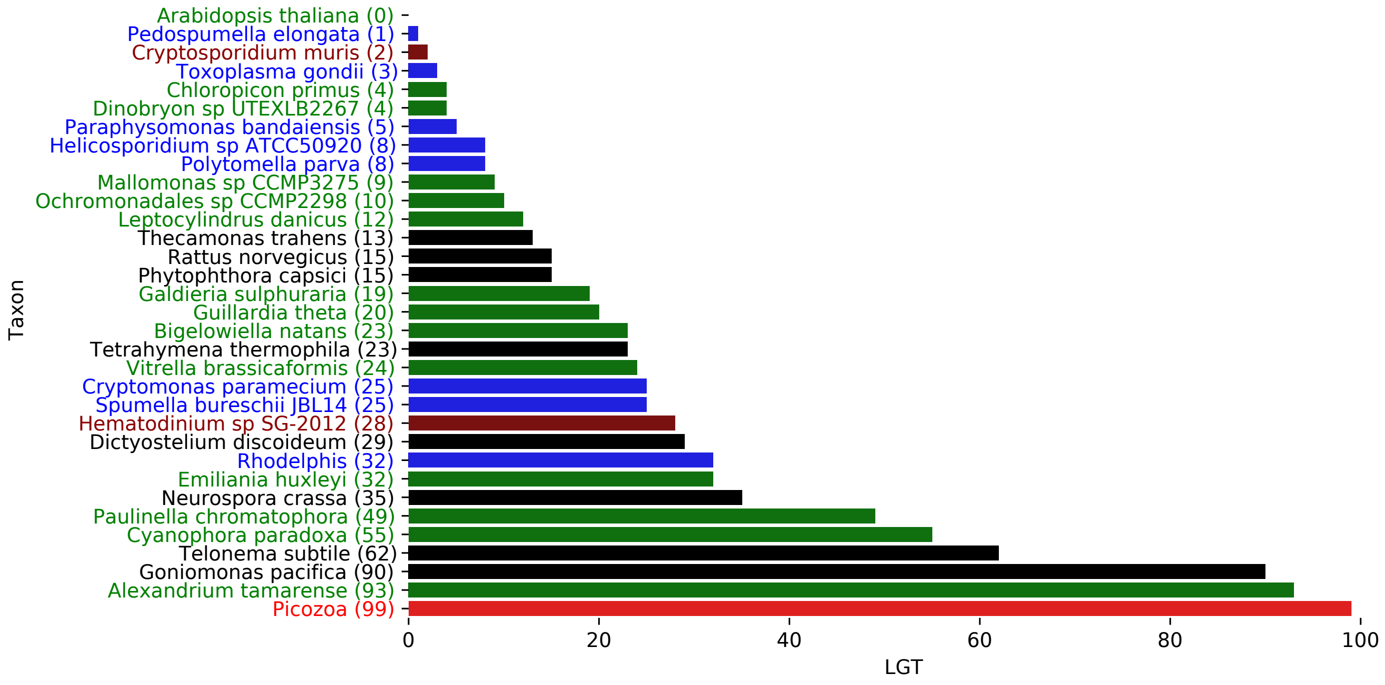

**Supplementary Figure 10: Number of inferred lateral gene transfers (LGT) across a selection of 33 species.** The species represent groups with a photosynthetic plastid (green), a non-photosynthetic plastid (blue), confirmed plastid loss (yellow) and no known plastid ancestry (black). These species serve as a comparison to Picozoa (orange).

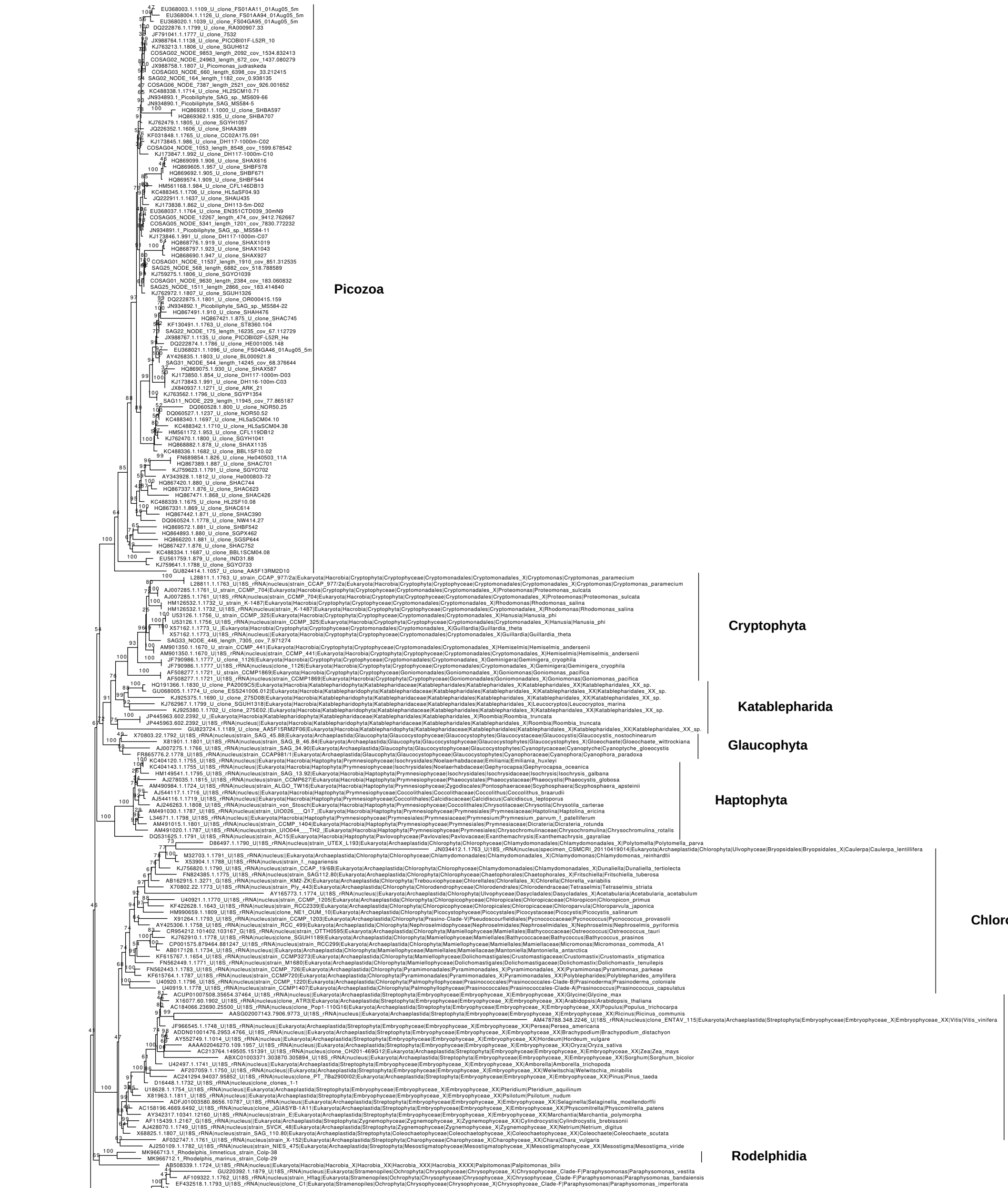

**Supplementary Figure 11: Maximum Likelihood tree of the 18S rRNA gene from the individual SAG assemblies and an extended number of reference sequences from Picozoa and other major eukaryotic groups from the PR2 database.** The tree was reconstructed under the model GTR+R5+F.

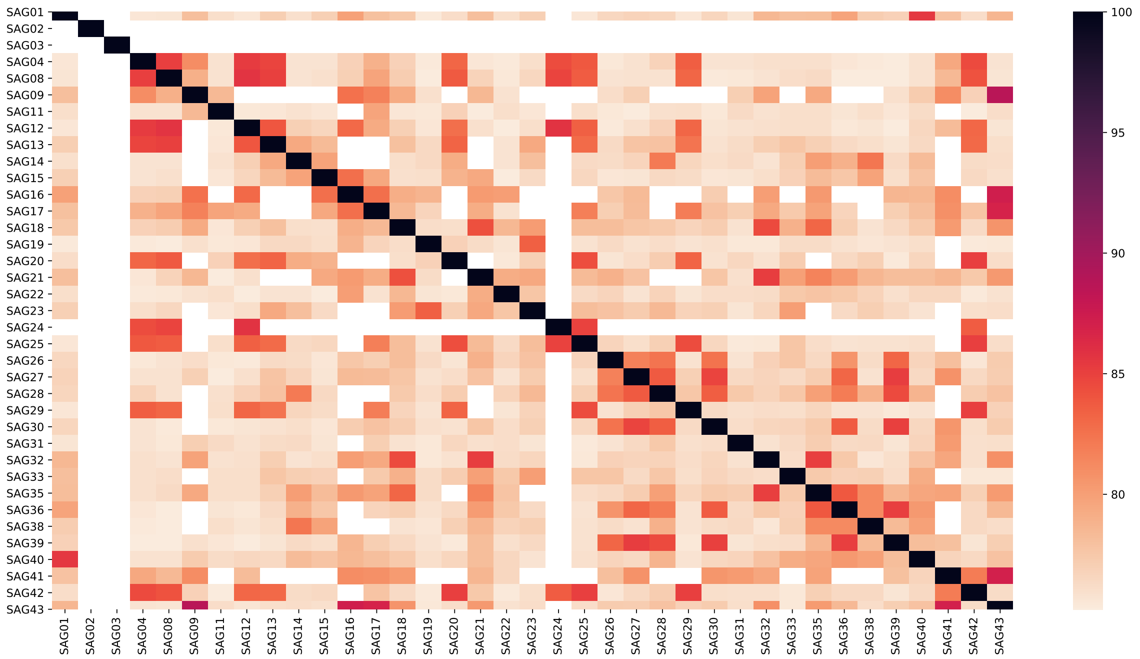

**Supplementary Figure 12: Heatmap showing pairwise ANI for 43 initial picozoan SAGs as estimated with FastANI.** Due to the incompleteness of the SAGs, many ANI values are zero, since there is no overlap between the assemblies.

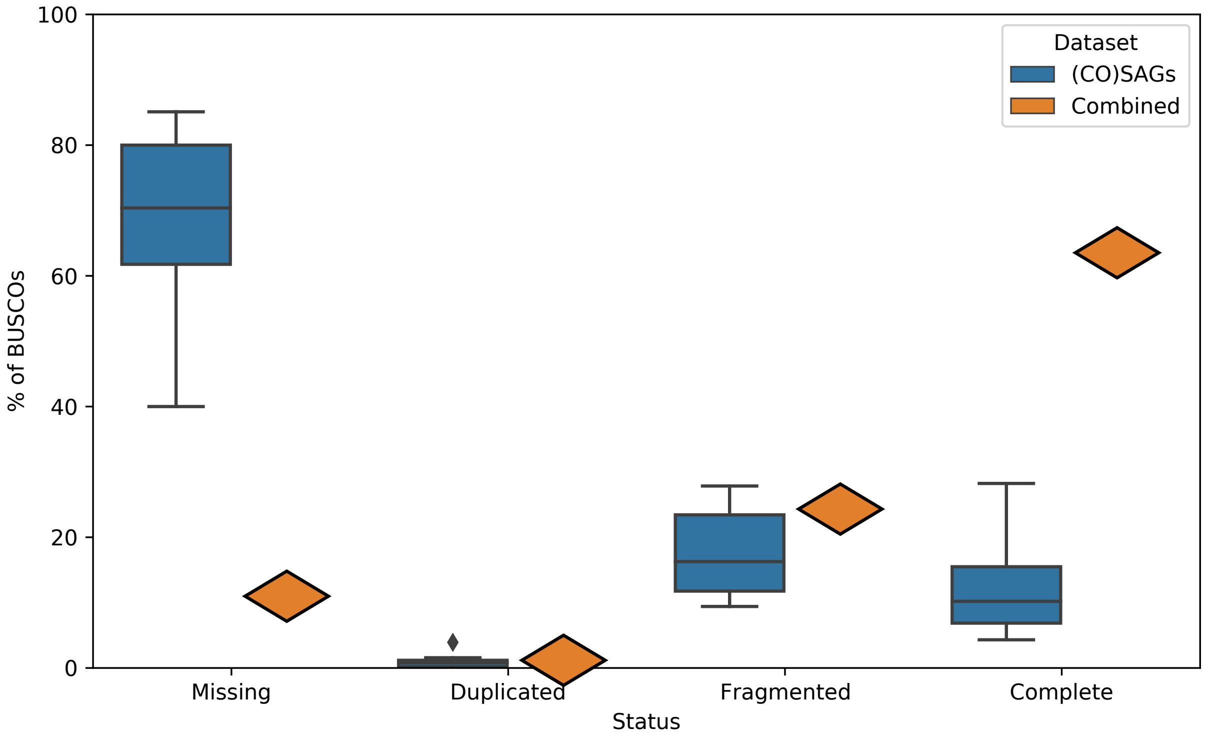

**Supplementary Figure 13: Boxplots of different BUSCO categories (Missing, Complete, Fragmented and Duplicated) for all selected SAGs/Co-SAGs.** Only SAGs that were used for Phylogenomic reconstruction were considered as well as the results for all these 10 assemblies combined. For the combined value, a BUSCO was considered complete if it was complete in at least one assembly.

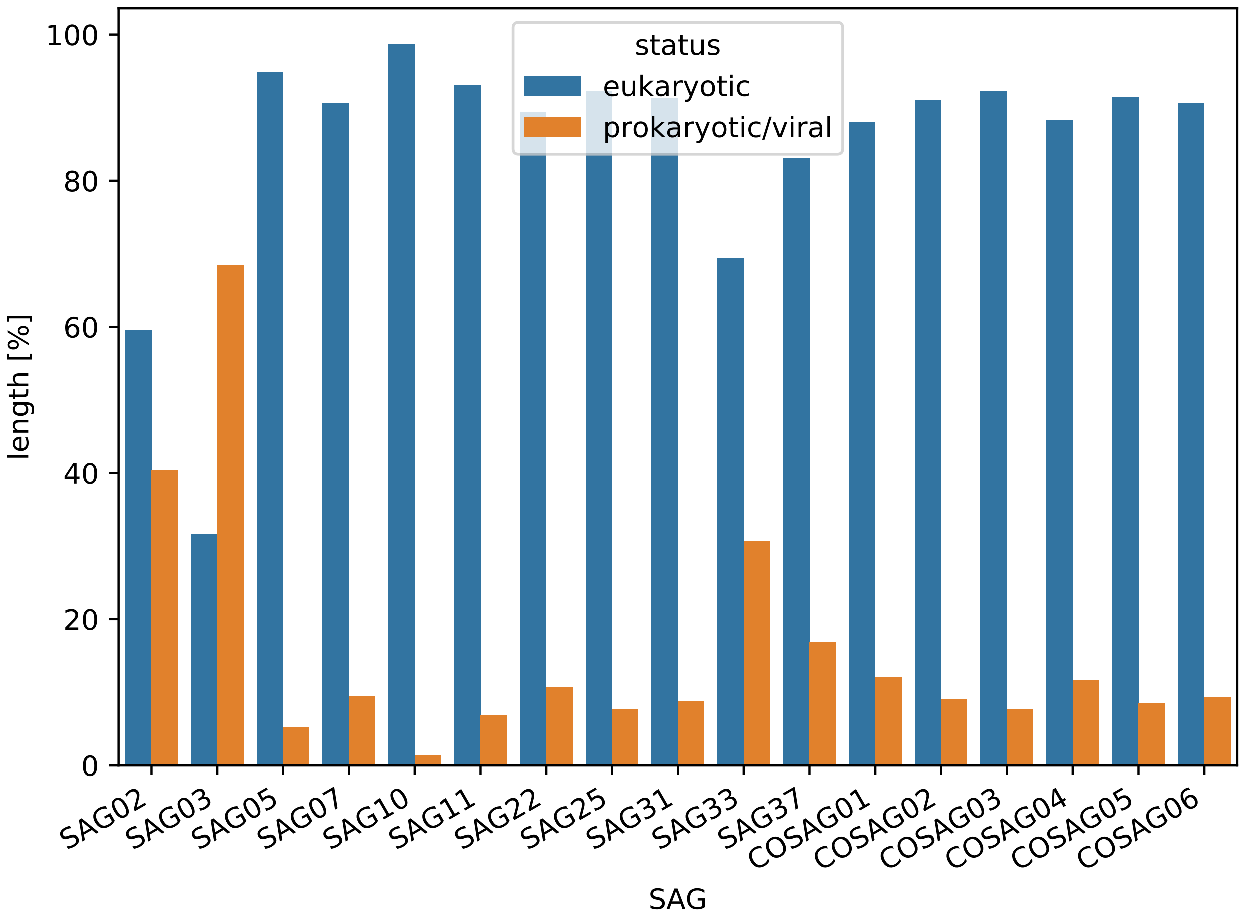

**Supplementary Figure 14: Contamination estimate for each of the 17 final SAGs/Co-SAGs.** All proteins were subjected to a DIAMOND blastp search against the ncbi NR database. If 60% of all proteins predicted from a contig only showed significant hits to prokaryotes or viruses, the contig was considered a putative contamination. Values correspond to fractions of contaminated/clean contigs of the total assembly length.

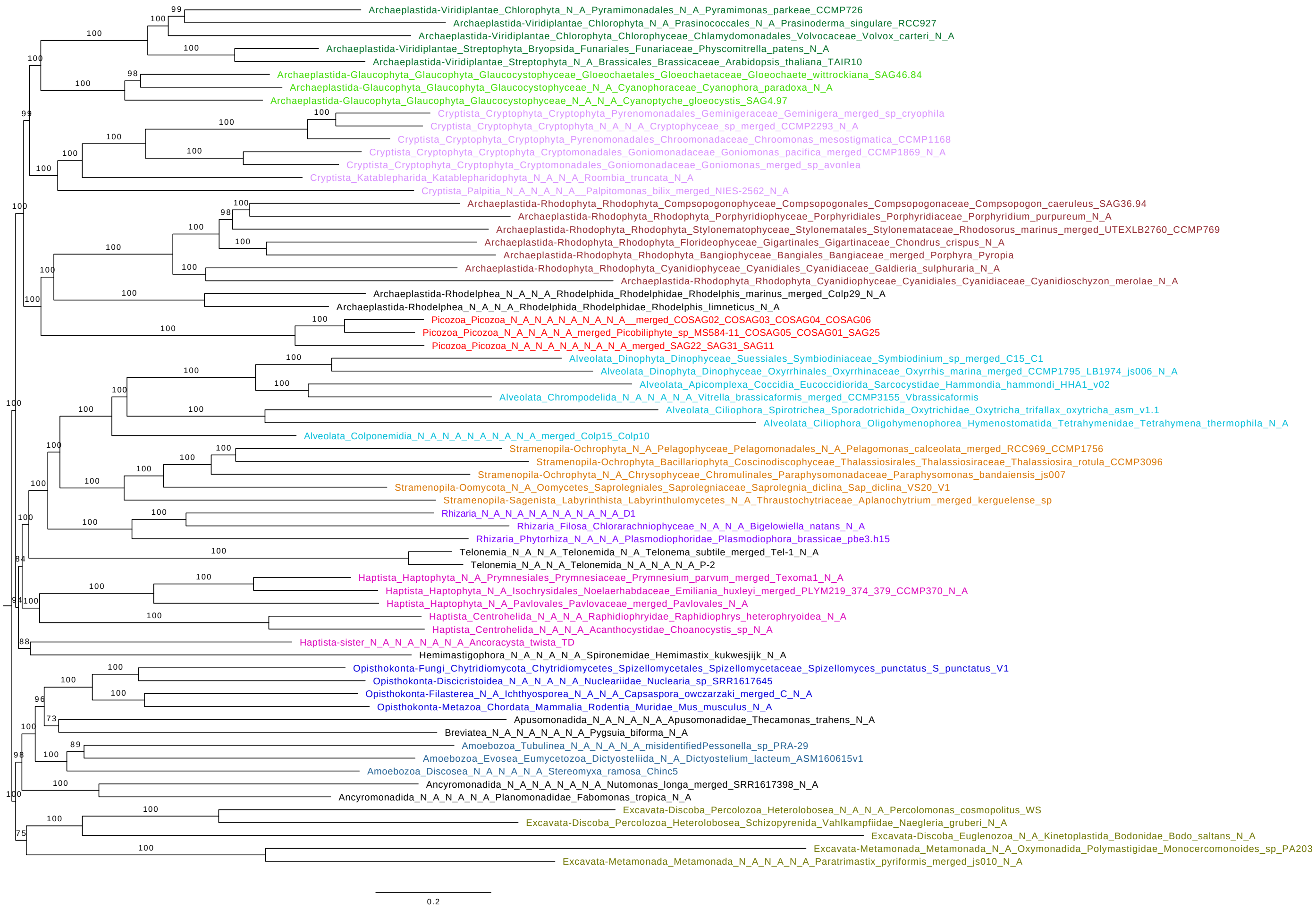

**Supplementary Figure 15: Maximum likelihood tree of eukaryotic species showing the position of Picozoa.** The tree is based on the concatenated alignment of 317 marker genes (filtered with Divvier) and was reconstructed using the PMSF approximation of the site-heterogeneous model LG+C60+F+G. Support values correspond to 100 non-parametric bootstrap replicates.

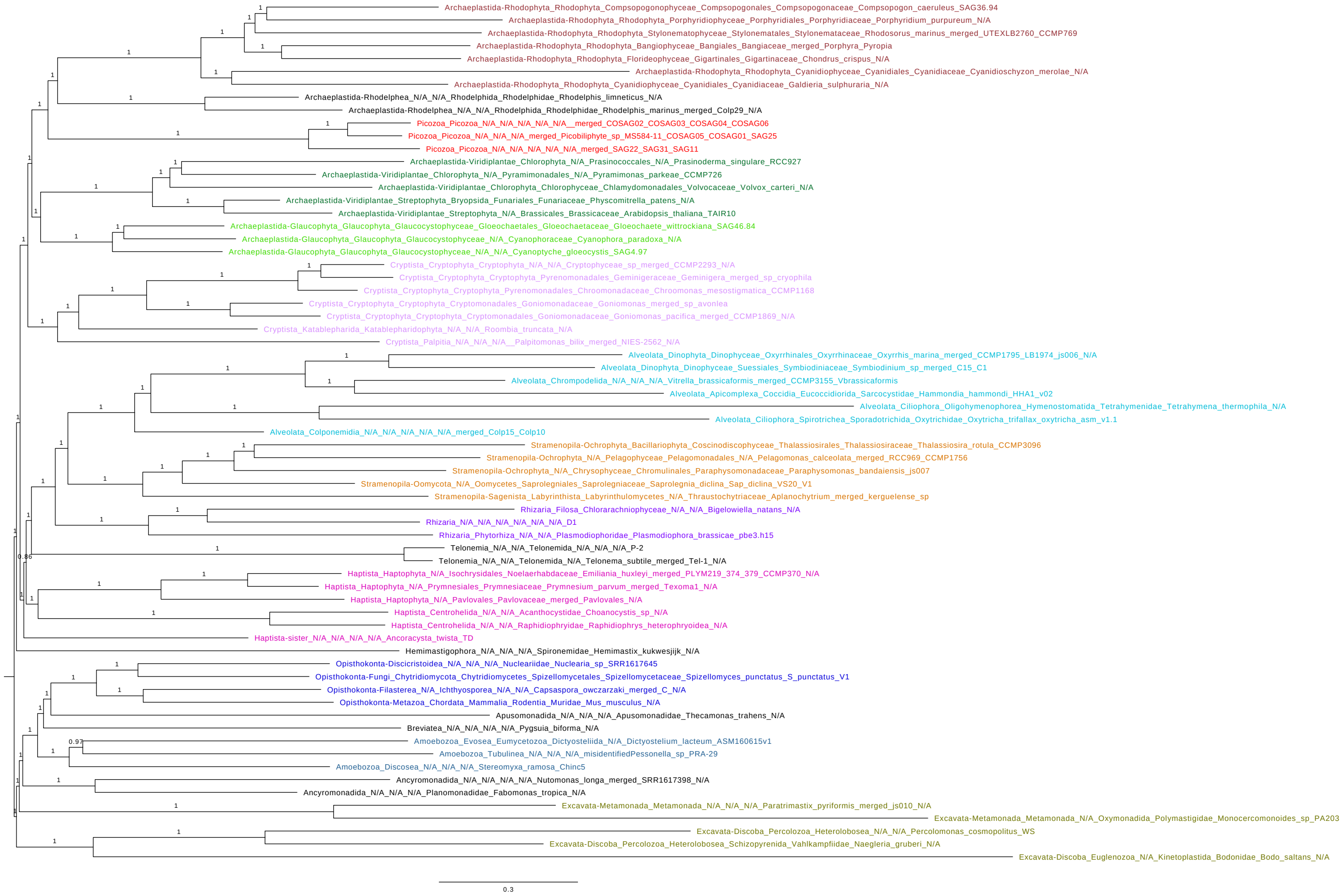

**Supplementary Figure 16: Bayesian phylogenetic tree made using PhyloBayes.** The tree is based on the concatenated alignment of 317 marker genes (filtered with BMGE). Shown is the consensus of chains 1 and 2 (maxdiff 0.26) after 3600 cycles with 1500 cycles removed as burn-in. Support values correspond to posterior probability values.

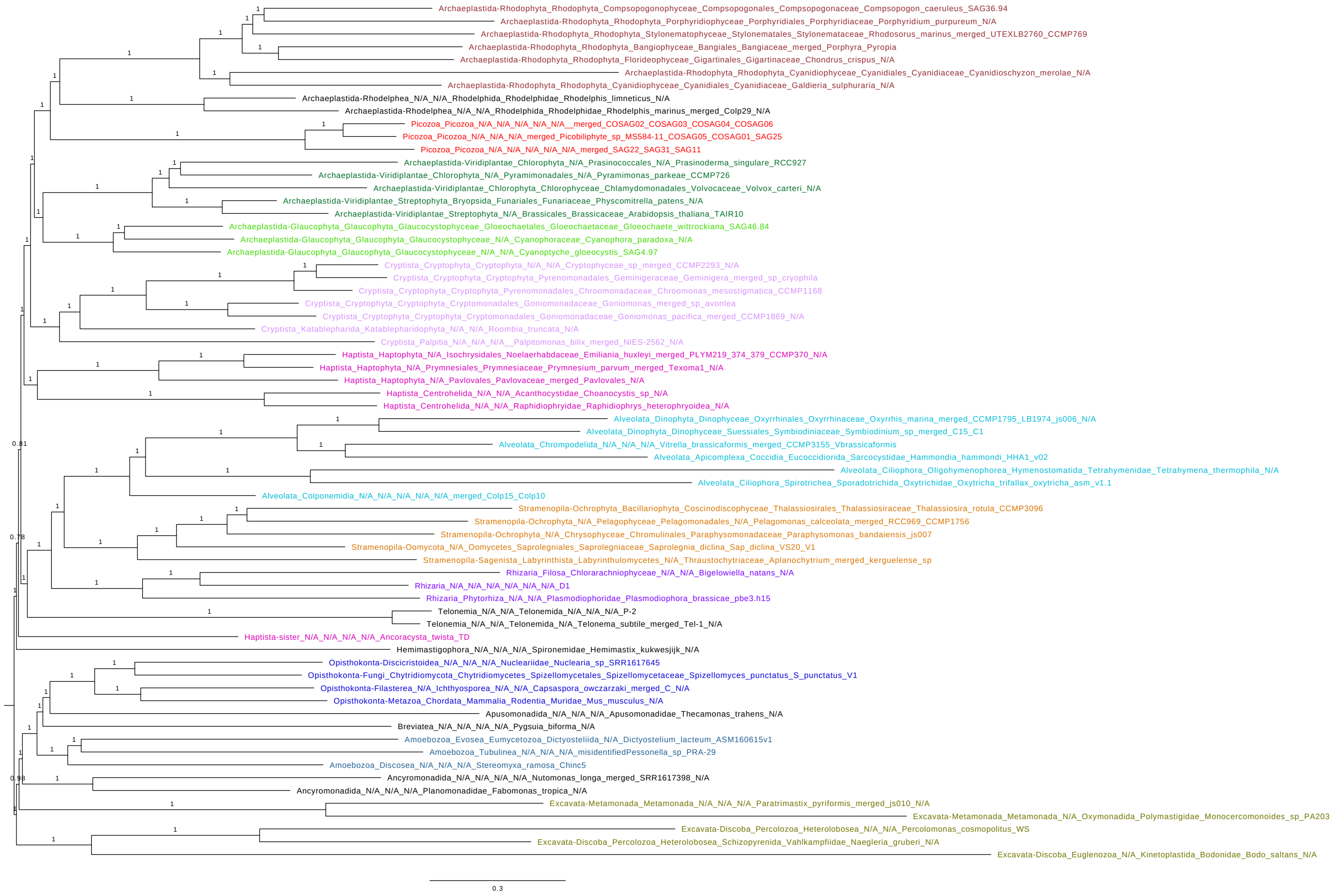

**Supplementary Figure 17: Bayesian phylogenetic tree made using PhyloBayes.** The tree is based on the concatenated alignment of 317 marker genes (filtered with BMGE). Shown is the consensus of chain 3 after 3600 cycles with 1500 cycles removed as burn-in. Support values correspond to posterior probability values.

**Supplementary Table 1:** Assembly/genome characteristics SAGs and CO-SAGs

**Supplementary Table 2:** Additional genomes added to the Phylogenomic dataset

**Supplementary Table 3:** Phylogenomic dataset taxon selection and taxon merging

**Supplementary Table 4:** Results from the AU and other topology tests performed with IQ-TREE

**Supplementary Table 5:** The Picozoan MS584-11 mitochondrial genome was used as a BLAST query into the 43 picozoan SAGs. Putative mitochondrial contigs were retrieved and used as queries into the NCBI nr database. If the top hit retrieved was MG202007.1, we considered the query sequence to be a bona fide mitochondrial contig.

**Supplementary Table 6:** Commonly retained plastid pathways and proteins

**Supplementary Table 7:** EGT clustering dataset taxon selection

**Supplementary Table 8:** Phylogenomic dataset taxon selection and taxon merging

**Supplementary Table 9:** EGT and LGT results for 33 selected species/groups

**Supplementary Data 1:** Untrimmed EEF2 alignment showing 2 amino acid signature in Rhodelphis, Rhodophyte, Chloroplastida, Cryptista, Haptophytes (but neither centrohelids not Ancoracysta) and Picozoa
